# The role and robustness of the Gini coefficient as an unbiased tool for the selection of Gini genes for normalising expression profiling data

**DOI:** 10.1101/718007

**Authors:** Marina Wright Muelas, Farah Mughal, Steve O’Hagan, Philip J. Day, Douglas B. Kell

## Abstract

We recently introduced the Gini coefficient (GC) for assessing the expression variation of a particular gene in a dataset, as a means of selecting improved reference genes over the cohort (‘housekeeping genes’) typically used for normalisation in expression profiling studies. Those genes (transcripts) that we determined to be useable as reference genes differed greatly from previous suggestions based on hypothesis-driven approaches. A limitation of this initial study is that a single (albeit large) dataset was employed for both tissues and cell lines.

We here extend this analysis to encompass seven other large datasets. Although their absolute values differ a little, the Gini values and median expression levels of the various genes are well correlated with each other between the various cell line datasets, implying that our original choice of the more ubiquitously expressed low-Gini-coefficient genes was indeed sound. In tissues, the Gini values and median expression levels of genes showed a greater variation, with the GC of genes changing with the number and types of tissues in the data sets. In all data sets, regardless of whether this was derived from tissues or cell lines, we also show that the GC is a robust measure of gene expression stability. Using the GC as a measure of expression stability we illustrate its utility to find tissue- and cell line-optimised housekeeping genes without any prior bias, that again include only a small number of previously reported housekeeping genes. We also independently confirmed this experimentally using RT-qPCR with 40 candidate GC genes in a panel of 10 cell lines. These were termed the Gini Genes.

In many cases, the variation in the expression levels of classical reference genes is really quite huge (e.g. 44 fold for GAPDH in one data set), suggesting that the cure (of using them as normalising genes) may in some cases be worse than the disease (of not doing so). We recommend the present data-driven approach for the selection of reference genes by using the easy-to-calculate and robust GC.

## Background

In a recent paper [1], we introduced the Gini index (or Gini coefficient, GC) [2-5] as a very useful, nonparametric statistical measure for identifying those genes whose expression varied least across a large set of samples (when normalised appropriately [6] to the total expression level of transcripts). The GC is a measure that is widely used in economics (e.g. [4, 7-12]) to describe the (in)equality of the distribution of wealth or income between individuals in a population. However, although it could clearly be used to describe the variation in any other property between individual examples [13-16]), it has only occasionally been used in biochemistry [1, 5, 17-22]. Its visualisation and calculation are comparatively straightforward (Fig 1): individual examples are ranked on the abscissa in increasing order of the size of their contribution, and the cumulative contribution is plotted against this on the ordinate. The GC is given by the fractional area mapped out by the resulting ‘Lorenz’ curve (Fig 1). For a purely ‘socialist’ system in which all contributions are equal (GC = 0), the curve joins the normalised 0,0 and 1,1 axes, while for a complete ‘autocracy’, in which the resource or expression is held or manifest by only a single individual (GC=1), the ‘curve’ follows the two axes (0,0 → 1,0 → 1,1).

**Fig 1.**
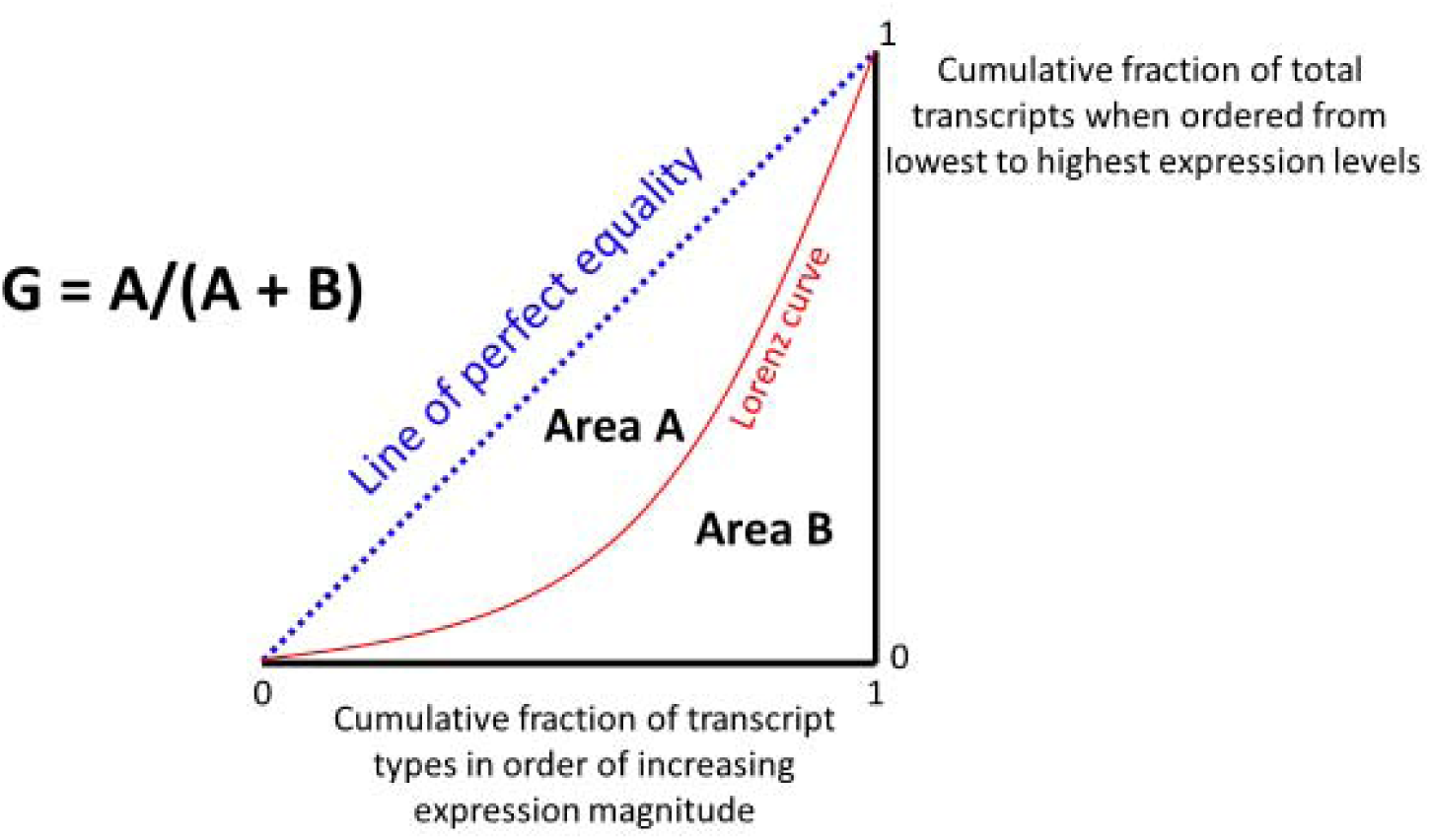
Graphical indication of the means by which we calculate the Gini coefficient.

Since the early origins of large-scale nucleic acid expression profiling, especially those using microarrays [23-25], it has been clear that expression profiling methods are susceptible to a variety of more or less systematic artefacts <u>within an experiment</u>, whose resolution would require or benefit from some kind of normalisation (e.g. [26-36]). By this (‘normalisation of the first kind’), and what is typically done, we mean the smoothing out of genuine artefacts <u>within</u> an arrray or a run, that occur simply due to differences in temperature or melting temperature or dye binding or hybridisation and cross-hybridisation efficiency (and so on) across the surface of the array. This process can in principle use reference genes, but usually exploits smoothing methods that normalise geographically local subsets of the genes to a presumed distribution.

Even after this is done, there is a second level of normalisation, that <u>between</u> chips or experiments, that is usually done separately, not least because it is typically much larger and more systematic, especially because of variations in the total amount of material in the sample analysed or of the overall sensitivity of the detector (much as is true of the within-run versus between-run variations observed in mass spectrometry experiments [37, 38]). This kind of normalising always requires ‘reference’ genes whose expression varies as little as possible in response to <u>any</u> changes in experimental conditions. The same is true for expression profiling as performed by qPCR [39-44], where the situation is more acute regarding the choice of reference genes since primers must be selected for these *a priori*. Commonly, the geometric mean of the expression levels of that or those that vary the least is selected as the ‘reference’. The question then arises as to which are the premium ‘reference’ genes to choose.

Perhaps surprisingly [45], rather than simply letting the data speak for themselves, choices of candidate reference genes were often made on the basis that reference genes should be ‘housekeeping’ genes that would simply be assumed (‘hypothesised’) to vary comparatively little between cells, be involved in nominal routine metabolism and also that they should have a reasonably high expression level (e.g. [46-63]). This is not necessarily the best strategy, and there is in fact (and see below) quite a wide degree of variation of the expression of most standard housekeeping genes between cells or tissues (e.g. [50, 59, 62, 64-76]). Indeed, Lee et al [66] stated explicitly that housekeeping genes may be uniformly expressed in certain cell types but may vary in others, especially in clinical samples associated with disease.

It became obvious that an analysis of the GC of the various genes was actually precisely what was required to assess those ‘housekeeping’ (or any other) genes that varied least across a set of expression profiles, and we found 35 transcripts for which the GC was 0.15 or below when assessing 56 mammalian cell lines taken from a wide variety of tissues [1]. These we refer to as the ‘Gini genes’. Most of these were ‘novel’ as they had never previously been considered as reference genes, and we noted that their Gini indices were significantly smaller (they were more stably expressed) than were those of the more commonly used reference genes [63]. However, this analysis was done on only one (albeit large) dataset of gene expression profiles. While some of the compilations (e.g. [62, 77]) contain massive amounts of expression profiling data, many of these, especially the older ones, may well be of uncertain quality. Thus, especially since the GC is very prone to being raised by small numbers of large outliers, we decided for present purposes that we should compare our analyses of candidate Gini genes using a smaller but carefully chosen set of expression profiling experiments. The more modern RNA-seq (e.g. [78-82]), in which individual transcripts are simply counted digitally via direct sequencing, is seen as considerably more robust [78, 83, 84] and sensitive [85, 86], and so we selected additional large and recent datasets that used RNA-seq in cell lines and tissues (Table *1*). We note too that the precision of these digital methods (as with other, digital, single-molecule strategies [87-89]), means that the requirement for reasonably high-level expression levels is much less acute.

**Table 1.** Studies used for assessing proposed stable reference genes.

| Study short name | Comments | Reference |
| --- | --- | --- |
| GiniGene | Study presenting novel potential housekeeping genes in cells and tissues from the HPA project cell and tissue RNASeq data. | [1] |
| geNorm or Vandesompele | Classic set of reference genes in tissues and a means of analysing them | [63] |
| Eisenberg | Very detailed analysis of housekeeping/reference genes in tissues using the **Illumina Body Map study** of RNA-seq of 16 Human Tissues. E-MTAB-513. | [46] |
| Lee | Two novel reference genes from a detailed analysis of 281 **normal tissue samples from 17 different organs then compares between disease states m and cell lines.** | [131] |
| Caracausi | 646 expression profile data sets from 54 different human **tissues.** | [62] |

In a similar vein (Table 2), we selected a small number of reasonably detailed studies in which particular housekeeping genes had been proposed as reference genes.

**Table 2.** Studies used for expression profiling data.

| Dataset short name | Comments | Reference |
| --- | --- | --- |
| HPA | RNA-seq-based dataset from the Human Protein Atlas group. Two data sets available: one of 19,628 protein coding genes in 56 **cell lines** (HPA_C) and another of 19,613 protein coding genes in 59 **tissues** (HPA_T). | [90, 91, 138] |
| CCLE | RNA-seq-based dataset (Cancer Cell Line Encyclopedia) of 58,035 genes in 934 human cancer **cell lines** (downloaded from EBI Expression Atlas E-MTAB2770). | [139] |
| Klijn / Genentech | RNA-seq-based analysis of 57,711 genes in 622 human cancer **cell lines** (downloaded from EBI Expression Atlas E-MTAB-2706). | [140] |
| GTEx | RNA-Seq data of 46,711 genes in 53 human **tissue** samples from the | [141] |
|  | Genotype-Tissue Expression (GTEx) project (downloaded from EBI Expression Atlas E-MTAB-5214). |  |
| PCAWG | RNA-Seq of 46,816 genes in 76 **tissues**, cancer and normal, from The International Cancer Genome Project: Pan Cancer Analysis of Whole Genomes ((downloaded from EBI Expression Atlas E-MTAB-5200). | Unpublished, may be subject to publication embargo until July 25<sup>th</sup>, 2019<br>https://dcc.icgc.org/pcawg |
| HBM | Illumina Body Map: RNA-seq of 16 Human **Tissues**. E-MTAB-513. Used by Eisenberg and colleagues in their analysis of housekeeping/ reference genes in **tissues**. | [46] |

To our knowledge, there are no large-scale studies to determine housekeeping genes in large, cell-line cohorts; the present paper serves to provide one. In addition, we include an experimental RT-qPCR analysis of a subset of the Gini genes.

## Results

### The Gini Coefficient as a robust measure of gene expression stability in multiple cell-line data sets

We previously identified a number of genes in the Human Protein Atlas (HPA) cell line data set [90] with very low expression variability and thus potential for use as reference genes [91]. However, we did not compare these Gini genes to other genes that have previously been proposed as housekeeping genes. We therefore performed a similar analysis using the potential housekeeping genes we proposed in [91] as well as other reference genes proposed in other studies (Table 2) with additional large RNA-Seq cell line data sets (Table *1*).

Fig 2A shows a plot of the GC of a variety of candidate Gini genes versus their median expression level in the HPA cell lines dataset set [90]. It is clear that genes we identified previously have much lower GC values in the HPA dataset than do any of the others (just two, VPS29 and CHMP2A, were also identified by Eisenberg and Levenson and another, RPL41, by Caracausi). This is not at the expense of an unusually low expression (Fig 2A), a finding broadly confirmed when we look at the median expression levels for the CCLE dataset (Fig 2B) and of the Klijn dataset (Fig 2C).

**Fig 2.**
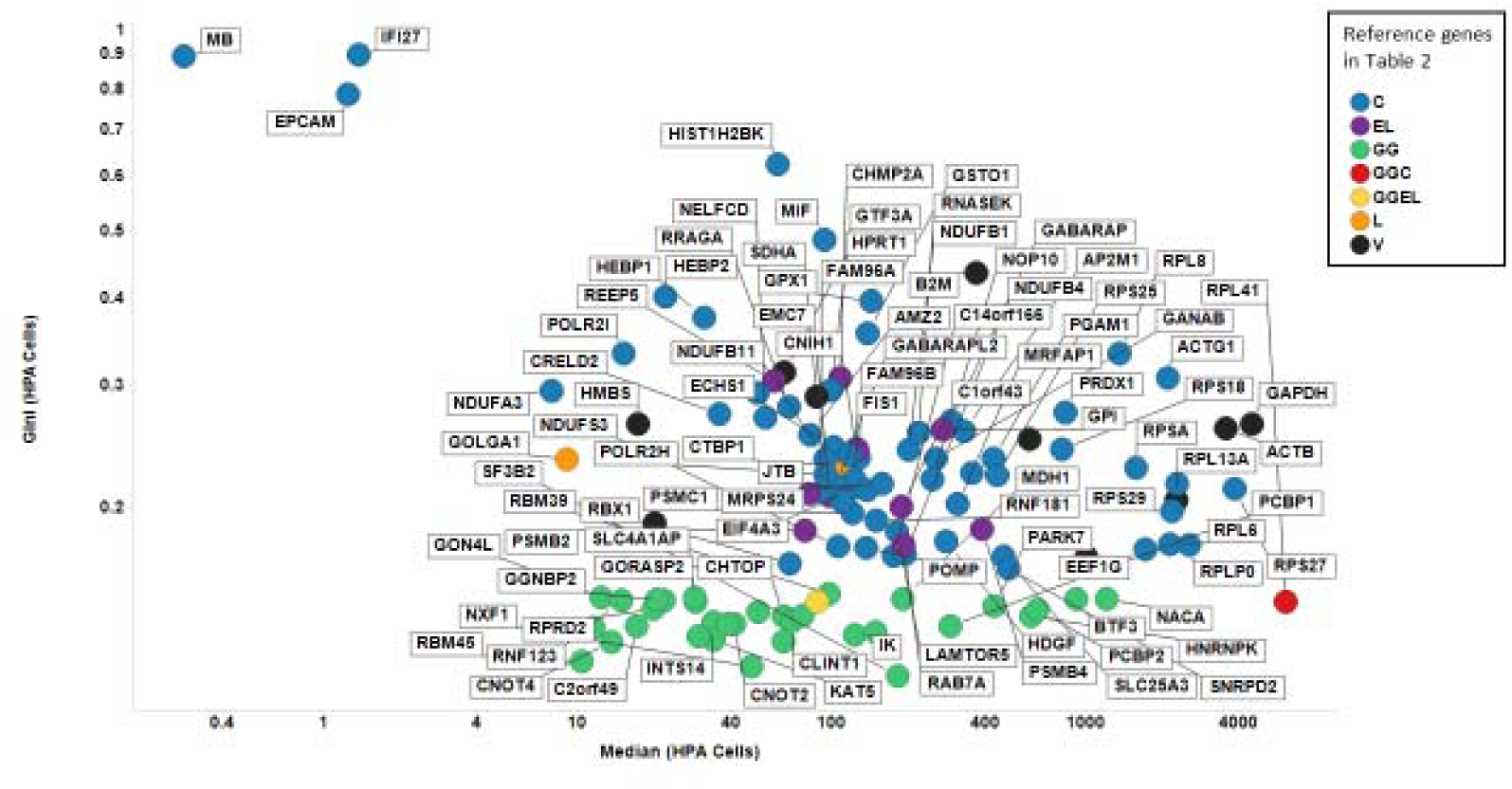
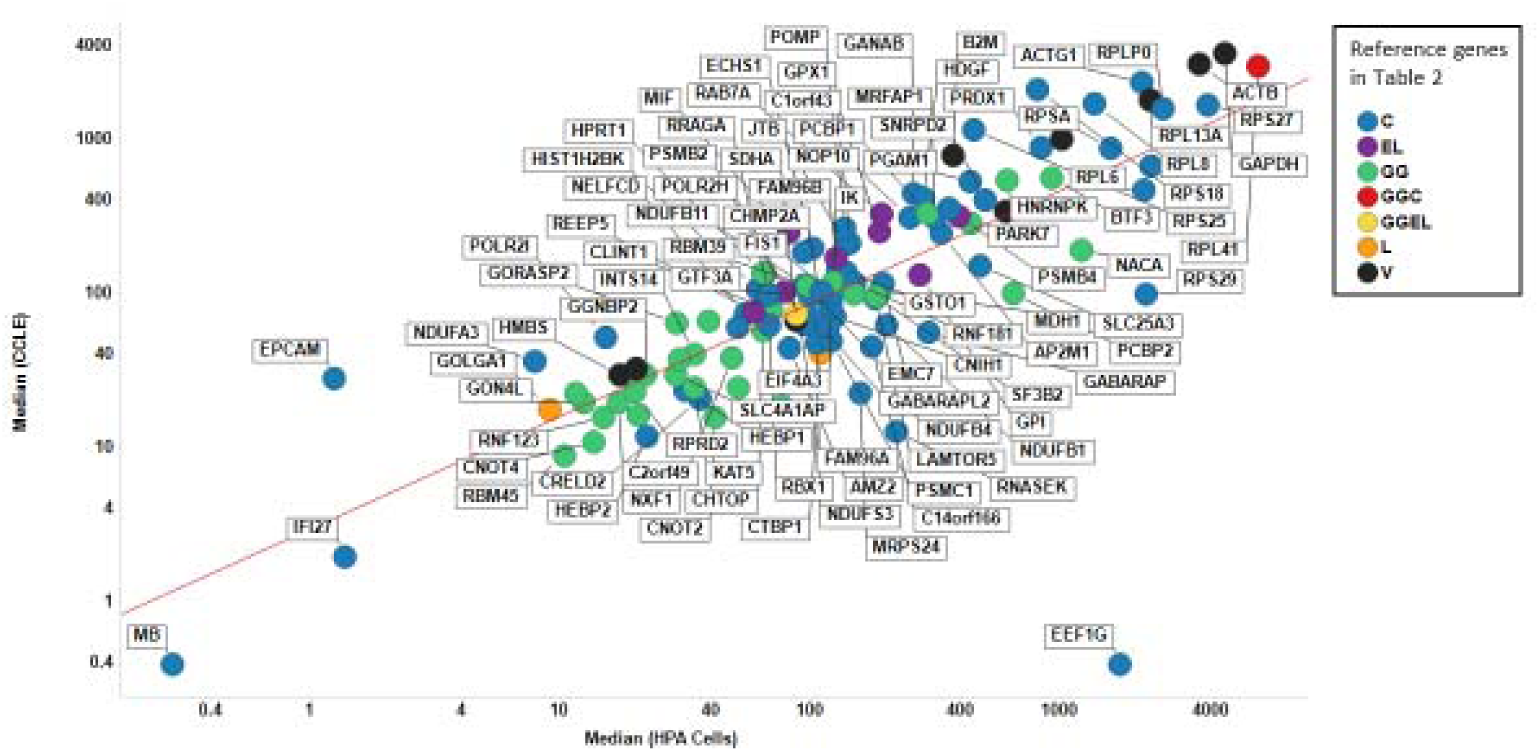
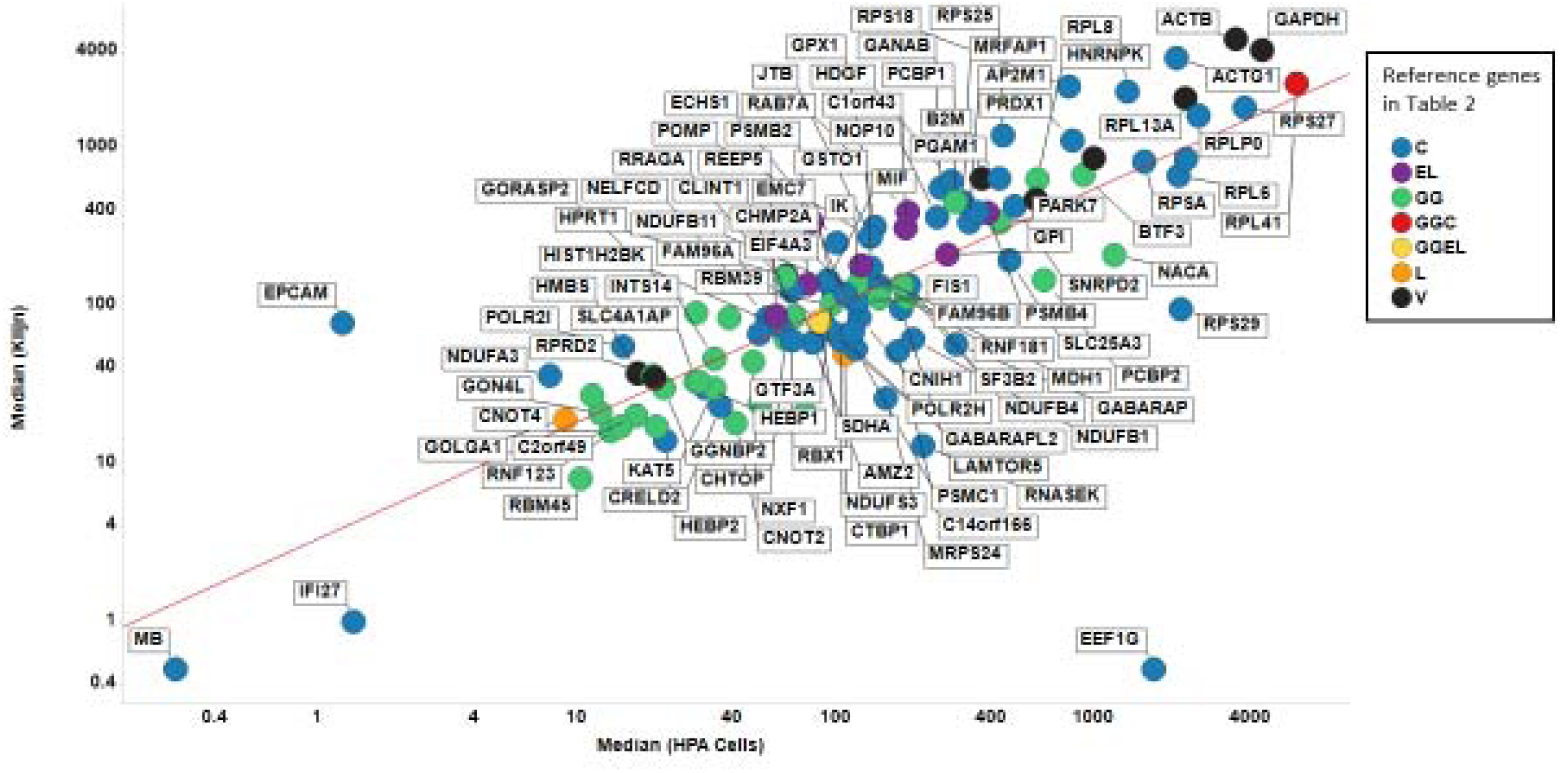
Gini coefficient and median expression levels of proposed reference genes in the HPA cell-line dataset. **A.** GC versus median expression level of HPA dataset. **B.** Median expression levels of CCLE vs HPA datasets. Line of best linear fit (in log space) shown is y = 0.991 + 0.827 X (r^2^ =0.606). **C.** Median expression levels of CCLE vs Klijn datasets. Line of best linear fit (in log space) shown is y = 0.998 + 0.804 X (r^2^ =0.593). Colour coding: red, GeneGini reference genes; blue Eisenberg & Levenson; yellow Vandesompele; green Lee; lilac both GeneGini and Eisenberg and Levenson.

Fig 3 shows the GC values for the various genes in two other datasets, viz CCLE and Klijn. Our previous Gini genes have a lower GC than that of any of the other housekeeping genes in 25 out of 38 cases in Klijn (all under 0.2) and in 26 out of 40 cases for CCLE (all under 0.22). In confirmation of this, and of the correlation found above between the median expression levels in CCLE and Klijn, the GC values are also well correlated with each other for the two datasets (Fig 3). Thus, although the absolute numbers are slightly larger than are those for the HPA dataset (unsurprisingly, given the much larger number of examples), the trend is still very clear: the GiniGenes remain the best among those variously proposed as reference genes in a variety of large and quite independent datasets. It also suggests that variations in the total amount of mRNA are not an issue either.

**Fig 3.**
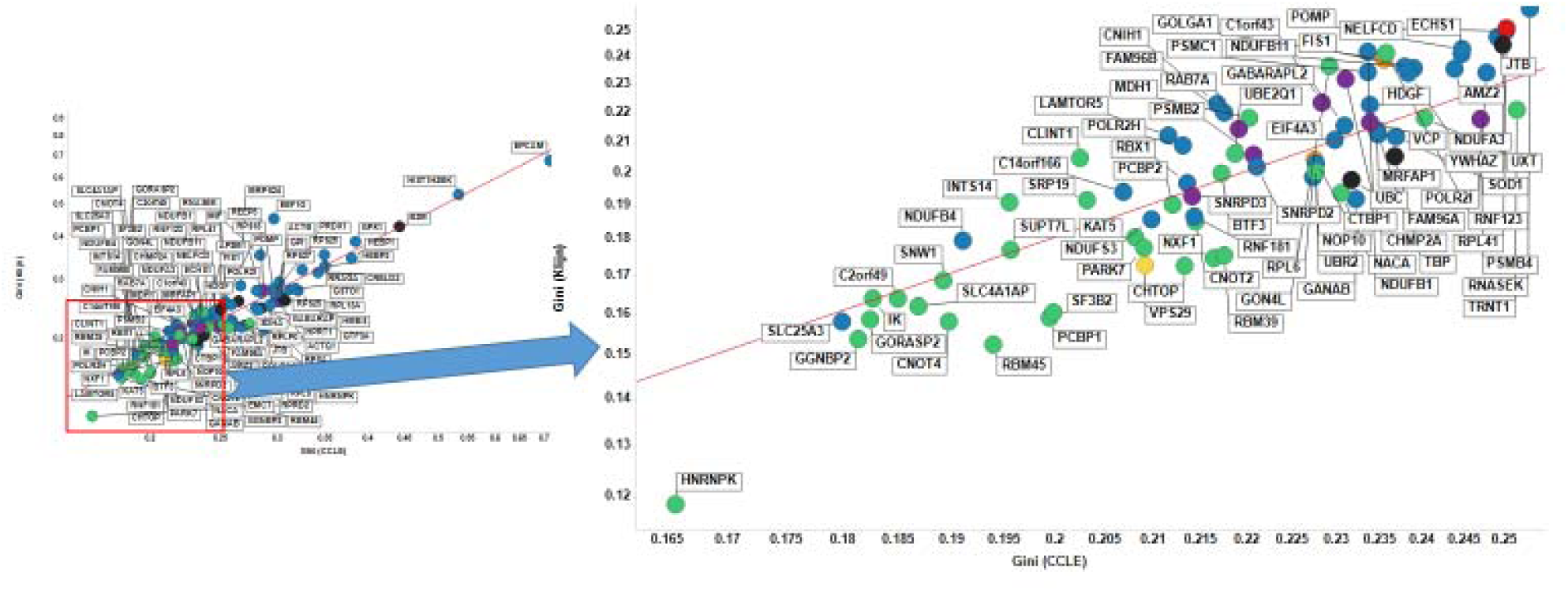
Gini coefficient of candidate reference genes in CCLE and Klijn/Genentech cell-line datasets. Left panel shows all proposed housekeeping genes considered in this study, with the right panel showing labels of those genes with a GC < 0.25. The line of best fit is y = −0.171 + 0.829x (r^2^ = 0.909). Colour code as in Fig 2.

Another common statistical measure, more resistant to individual outliers, is the interquartile ratio (the ratio between the 25^th^ and 75^th^ percentile when expression levels are ranked); by this measure too, the Gini genes that we uncovered previously stand out as being the least varying (Fig 4 A and B). This suggests that, as a measure of gene expression stability, the GC is robust: the GiniGenes have the lowest ratio between their maximum and minimum expression values in the HPA dataset (Fig 4C) and also the lowest interquartile ratio in their levels of expression in all three cell line data sets explored here (Fig. 4B and C) with good correlation between these two datasets.

**Fig 4.**
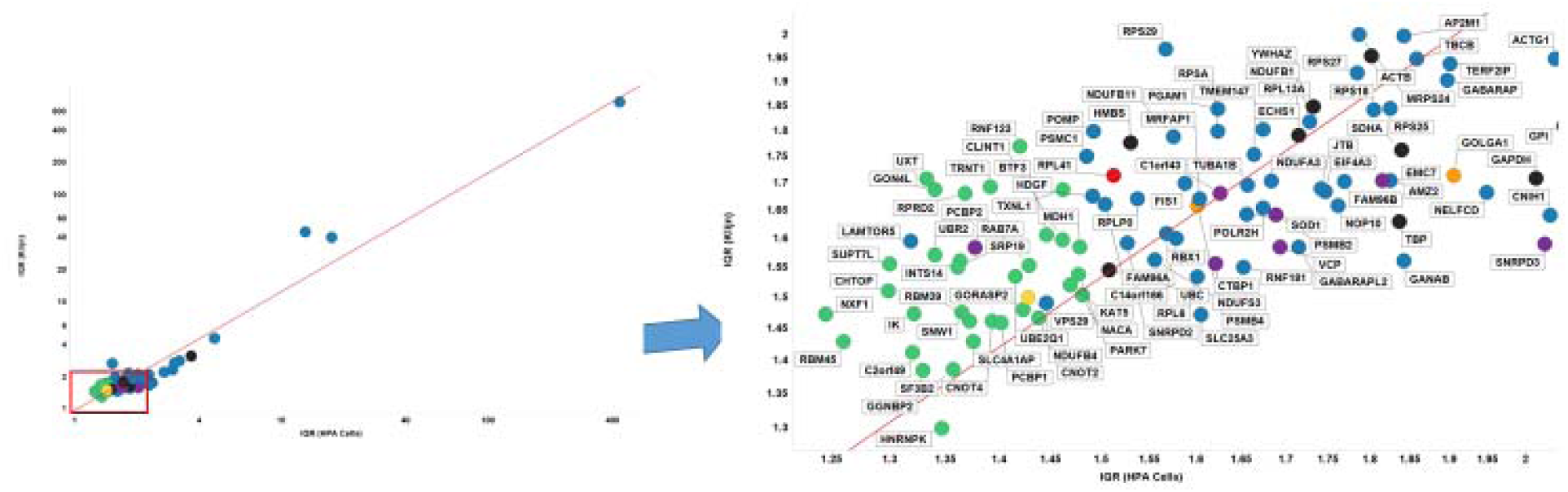
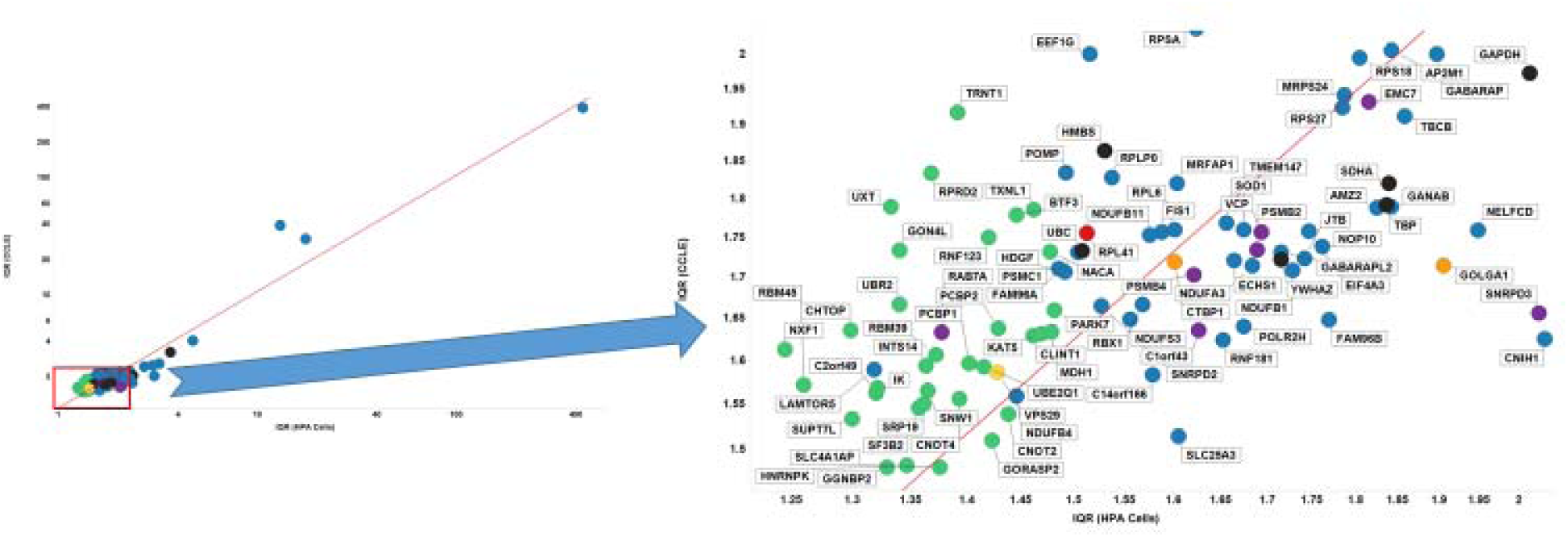
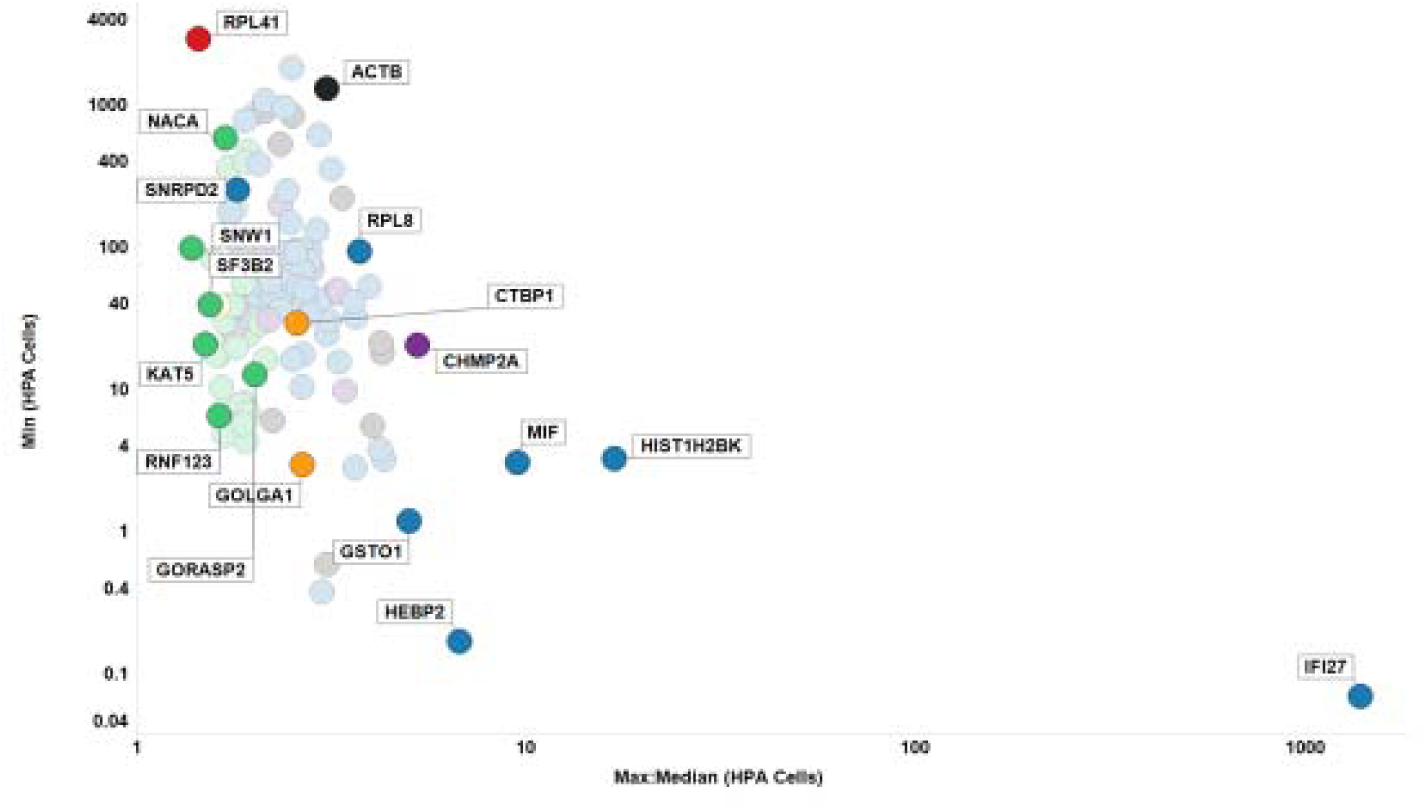
Robustness of the Gini coefficient. **A.** IQR of different genes in Klijn/Genentech vs HPA cell-line dataset. Left panel shows all genes considered in this study, with right panel showing genes with IQR < 2 in both datasets. Line of best linear fit (in log space) shown is y = 0.01 + 1.11 X (r^2^ =0.937) **B.** IQR of different genes in CCLE vs HPA cell-line dataset. Left panel shows all genes considered in this study, with right panel showing genes with IQR < 2 in both datasets. Line of best linear fit (in log space) shown is y = 0.04 + 0.99 X (r^2^ =0.930). **C.** Max:Mean vs Min expression levels in HPA data set. Colour code as in Fig 2.

### Use of the Gini Coefficient to find GiniGenes in an unbiased manner in cell-line data sets

Up to now, our analyses of these data sets have used a set of predefined genes to look at expression stability. We next sought to investigate whether the GC would highlight genes with high expression stability that have been reported by others or by ourselves when performing this analysis in a data-driven manner. To that end, we found 115 genes shared between the three data sets with a GC ≤ 0.2 (Fig. 5, 6). This value for the GC was chosen since reducing this to ≤ 0.15 meant no or very few genes were found in some data sets (e.g. no genes in the CCLE data set had a GC ≤ 0.15) and going above this meant the number of genes were unmanageable (e.g. 1051 genes with a GC ≤ 0.21 in the Klijn data set). Of the 115 genes shared between the datasets with GC <0.2, 13 were GiniGenes and two were housekeeping genes defined by Caracausi and colleagues (Fig. 5 B). When we selected the top 20 expressing genes in each data set, only 13 of these were common across these data sets; Table 3 shows some descriptive statistics of 13 of these, with descriptive statistics of all 115 genes found in Supplementary Table S1. Of these genes, two (HNRNPK and PCBP1) are GiniGenes and one (SLC25A3) is a gene previously reported by Caracausi et al. Seven out of the 13 genes (HNRNPK, HNRNPC, PCBPB, SF3B1, SRSF3, EDF1 and EIF4H) here share important roles in RNA transcription, translation and stability [92-100], are implicated in a number of diseases, including cancer [92, 95, 101-111], and some, such as SRSF3 are essential for embryo development [112]. Given their pivotal functions, it may be unsurprising that the expression of these genes are tightly regulated across cell lines of different tissue origins, even where these are cancer cell lines. Overall, the distribution, expression stability and important functional roles of these genes suggest that these are excellent housekeeping genes across different cell types.

**Fig 5.**
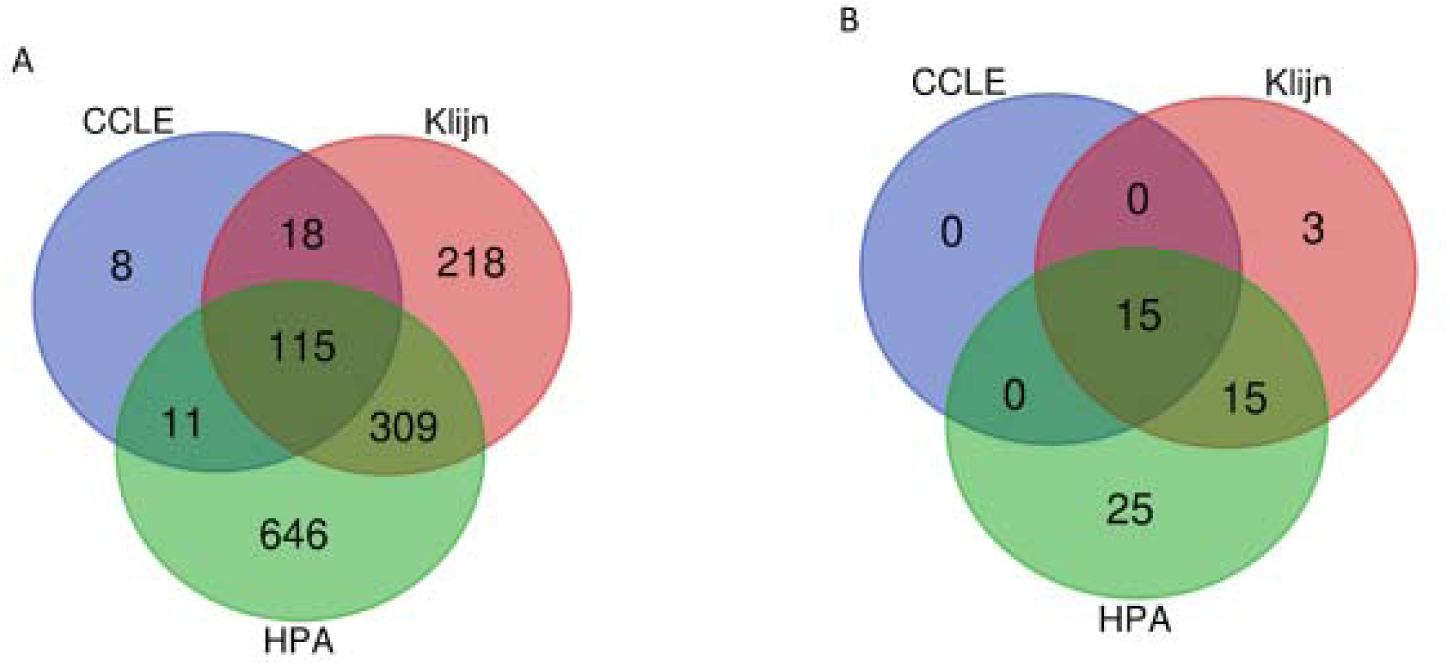
Shared and unique genes in HPA, CCLE and Klijn/Genentech cell-line data sets. **A.** Genes with a GC < 0.2 **B.** Housekeeping genes in Table 2 with GC < 0.2.

**Fig. 6.**
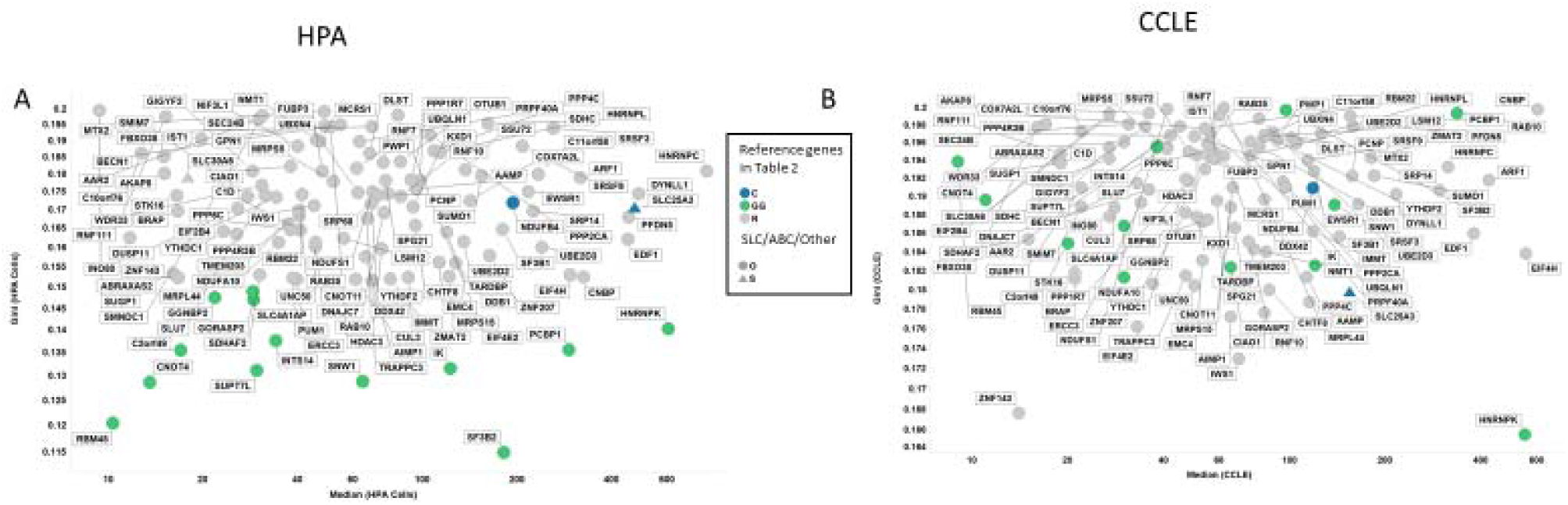
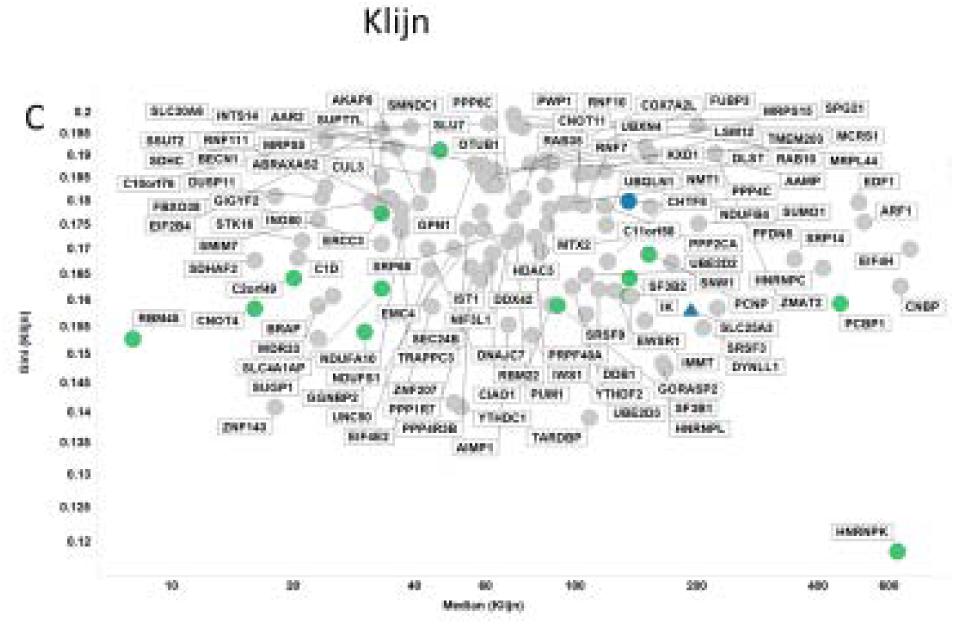
GC vs Median for 115 genes in **A.** HPA, **B.** CCLE and **C.** Klijn/Genentech cell-linedata sets. Colour coding: Blue, Caracausi; Green, GeneGini reference genes; Grey, neither. Shape coding: Circle, other; Triangle, SLC coding gene.

Of particular interest to us was finding one gene encoding a mitochondrial phosphate transporter protein (SLC25A3 [113]) to be within this list of the top expressing stably expressed genes. This might seem logical since mitochondrial ATP synthesis is required by all cell types and tissues.

Figure 7 shows the robustness of the GC for the subset of 115 genes common between the three data sets studied here with a low GC (<0.2). Lower Gini coefficients correlate with lower IQR and Max:median ratios (Fig7: only results for the Klijn data set are shown). The range of IQR values of these genes was smaller in the larger two data sets (CCLE, 1.42-1.67; Klijn, 1.30-1.64) than in the HPA data set (1.26-1.84) suggesting the measured expression values were more stable in the larger data sets (Supplementary Table S1). This may, however, be due to a larger number of cell lines in these two large datasets (934 and 622 in CCLE and Klijn) compared with the HPA data set (56 cell lines).

**Fig. 7.**
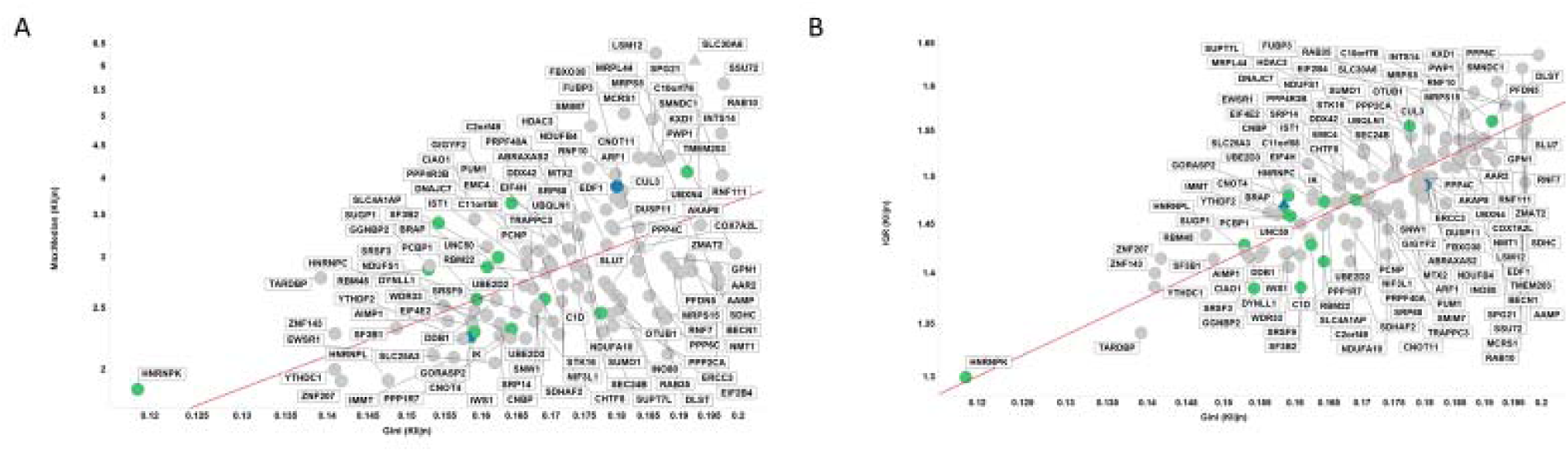
Robustness of GC for finding stably expressed genes using shared genes between HPA, CCLE and Klijn/Genentech cell-line data sets with GC < 0.2. Shown are the results for the Klijn/Genentech dataset **A.** IQR vs GC, **B.** Max:Mean vs Min. Colour coding: Blue, Caracausi; Green, GeneGini reference genes; Grey, neither. Shape coding: Circle, other; Triangle, SLC coding gene.

### Application of the Gini coefficient to human tissue RNA-Seq data sets

The results presented thus far are representative of human cell lines. Most reports in the literature regarding housekeeping genes refer to tissue expression data. This may be due to the cell lines being “dedifferentiated” with respect to the tissues from which they are derived [114].

In our previous report [1] we also analysed RNA-Seq data from tissues [90] and found 22 genes with a GC < 0.15, of which 3 (CHMP2A, VPS29 and PCBP1) were also found in cell line data with a GC <0.15. The median expression level and GC of these and other candidate GiniGenes in this tissue data set are shown in Figure 8. As with cell line data, the genes we previously identified (GGs, green dots in Fig 8) have much lower GCs in this tissue data set than do any of the other candidate GiniGenes, with only two of these genes (VPS29 and CHMP2A) identified previously by Eisenberg & Levenson [115]. The low GC value of these GiniGenes is not at the expense of low expression: of the 22 GiniGenes, 13 are expressed at a median level of between 40 and 200 TPM (see Supplementary Table S2). Moreover, the GC was also representative of the variation in expression of these genes (albeit influenced to a lesser extent by outliers), as shown in Fig. 9 A and B, with all GiniGenes having a GC < 0.15 and the lowest RSD (relative standard deviation), ranging from 24.096 % to 28.66 % and IQR (1.26 to 1.44) of this list of housekeeping genes. The expression of other housekeeping genes such as GAPDH, ACTB, RPL13A, SDHA, B2M was quite varied according to these measures. For example, the GC of GAPDH (a commonly used HKG) was 0.33, with a RSD of 72.4 % and IQR of 2.24, and for ACTB (another commonly used HKG) these values were 0.29, 55.24 %, and 2.11.

**Fig 8.**
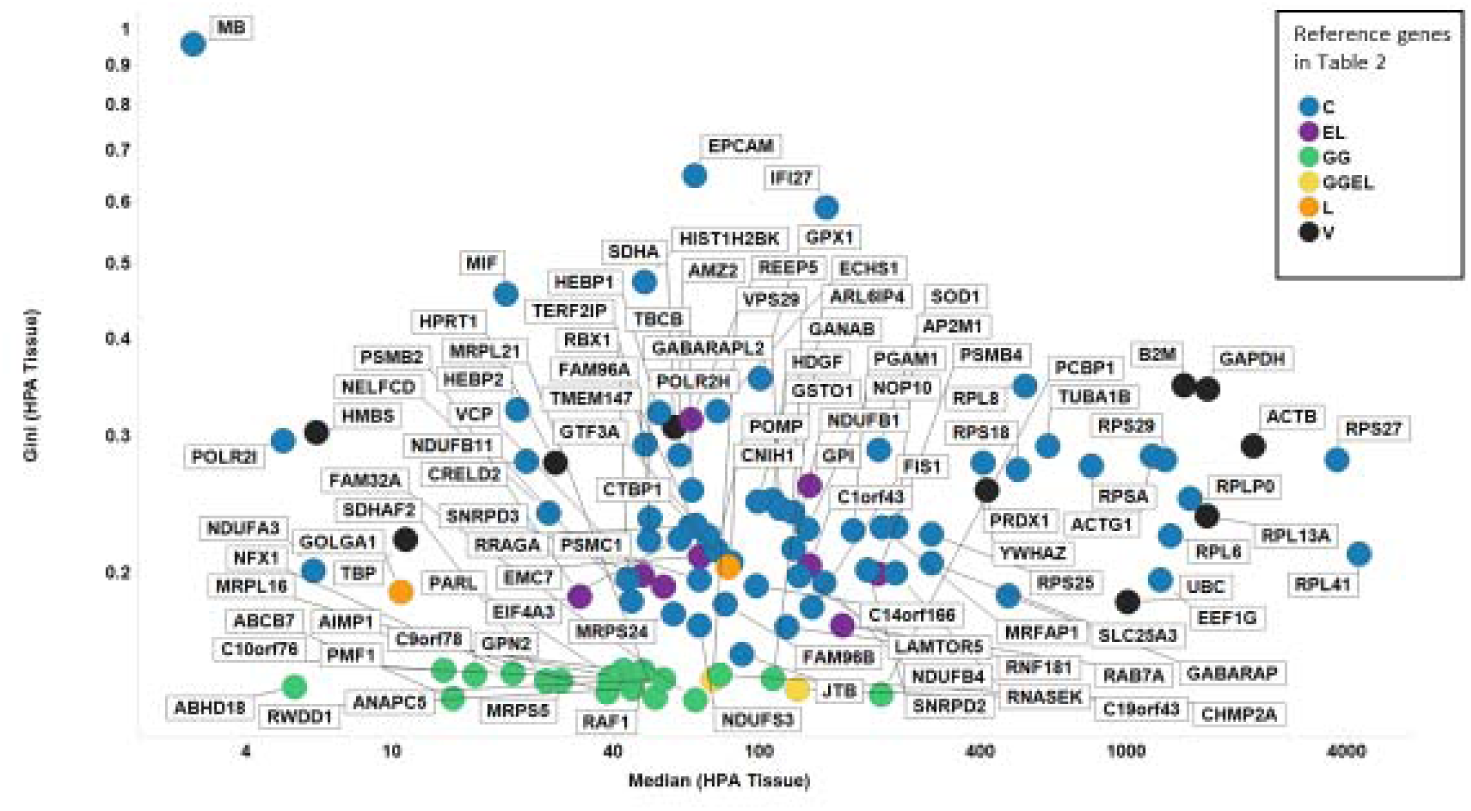
Gini coefficient and median expression levels of proposed reference genes in the HPA tissue dataset. Colour coding: blue, Caracausi; purple, Eisenberg and Levenson; green, GeneGini reference genes; yellow, both GeneGini and Eisenberg and Levenson; orange, Lee; black, Vandesompele.

**Fig 9.**
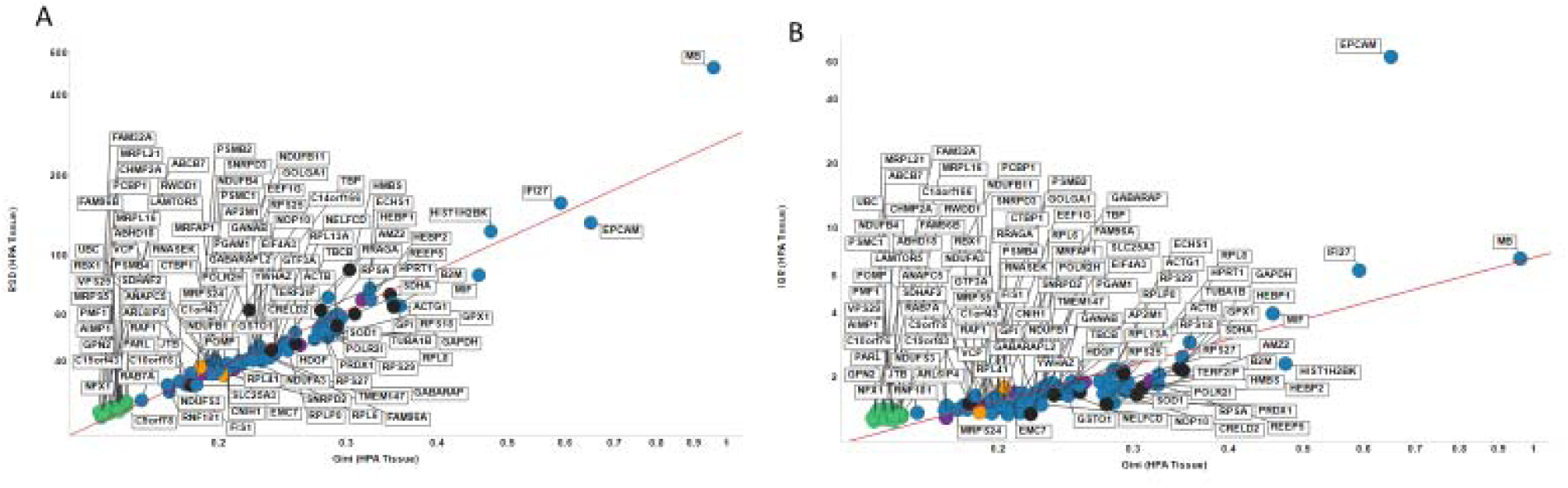
Robustness of the Gini coefficient in the HPA tissue data set. **A.** RSD versus Gini coefficient of candidate reference genes. Line of best linear fit (in log space) shown is y = 2.45 + 1.24 X (r^2^ =0.938) **B.** IQR versus Gini coefficient of candidate reference genes. Line of best linear fit (in log space) shown is y = 0.87 + 0.96 X (r^2^ =0.566). Colour code as in Fig 8.

The median expression levels of the proposed reference genes show a similar level of correlation between the data sets as was found with the cell line data (Figure S1 A-C), and GiniGenes displayed a mid-range level of expression. The GC of the tissue GiniGenes we proposed however, tended to be higher and more variable in their GC values than in the HPA dataset (Figure S2 A-C) suggesting that those genes may be representative of the HPA data set only. As an example, in the GTEx dataset only 28 genes had a GC < 0.2, of which the majority (17) were those reported by Caracausi and colleagues, and 7 were GiniGenes. The results here are likely influenced by the number and status (disease or normal) of the tissues analysed in the various data sets compared; for example, the GTEx data come from 53 different, normal human tissues, whereas the HPA tissue data include a mixture of disease and normal tissue samples. In addition, compared to the cell line data where hundreds (in the case of the Cancer Cell Line Encyclopedia) of cell lines were analysed, the number of tissues in these data sets was fewer than 100.

In the case of the data set used by Eisenberg and Levanon [115], viz. the Illumina Human Body Map (E-MTAB-513), 10 of the 11 housekeeping genes proposed here (which included 2 Gini Genes, CHMP2A and VPS29) had a GC ≤ 0.2 and were reasonably well expressed (with median expression levels between 50-270 TPM, see Supplementary Table S2 and Supplementary Fig S4). This may be compared to the 5 other GGs with GC < 0.2 in this data set whose expression value was lower, with median expression between 19-35 TPM. This suggests that finding suitable HKGs may be dependent on the data set itself, and the type of tissue under investigation.

We next sought to perform a more comprehensive and integrative analysis by filtering the tissue data sets to only include genes with a GC ≤ 0.2 to find common genes across these data sets with reasonable expression stability (Supplementary Table S3). As shown in Fig 10 only 15 genes were shared between the four data sets with a GC ≤ 0.2, none of which has been reported previously as a housekeeping genes. Table 4 shows some descriptive statistics of these genes. In any case, the names of the proteins encoded by these 15 genes suggest these play important and essential roles. The median expression values of these genes varied from around 10-450 TPM, with SNX3 (Sorting nexin-3 (Protein SDP3)) and COX4I1 (Cytochrome c oxidase subunit 4 isoform 1) being consistently the two highest-expressing genes.

**Fig 10.**
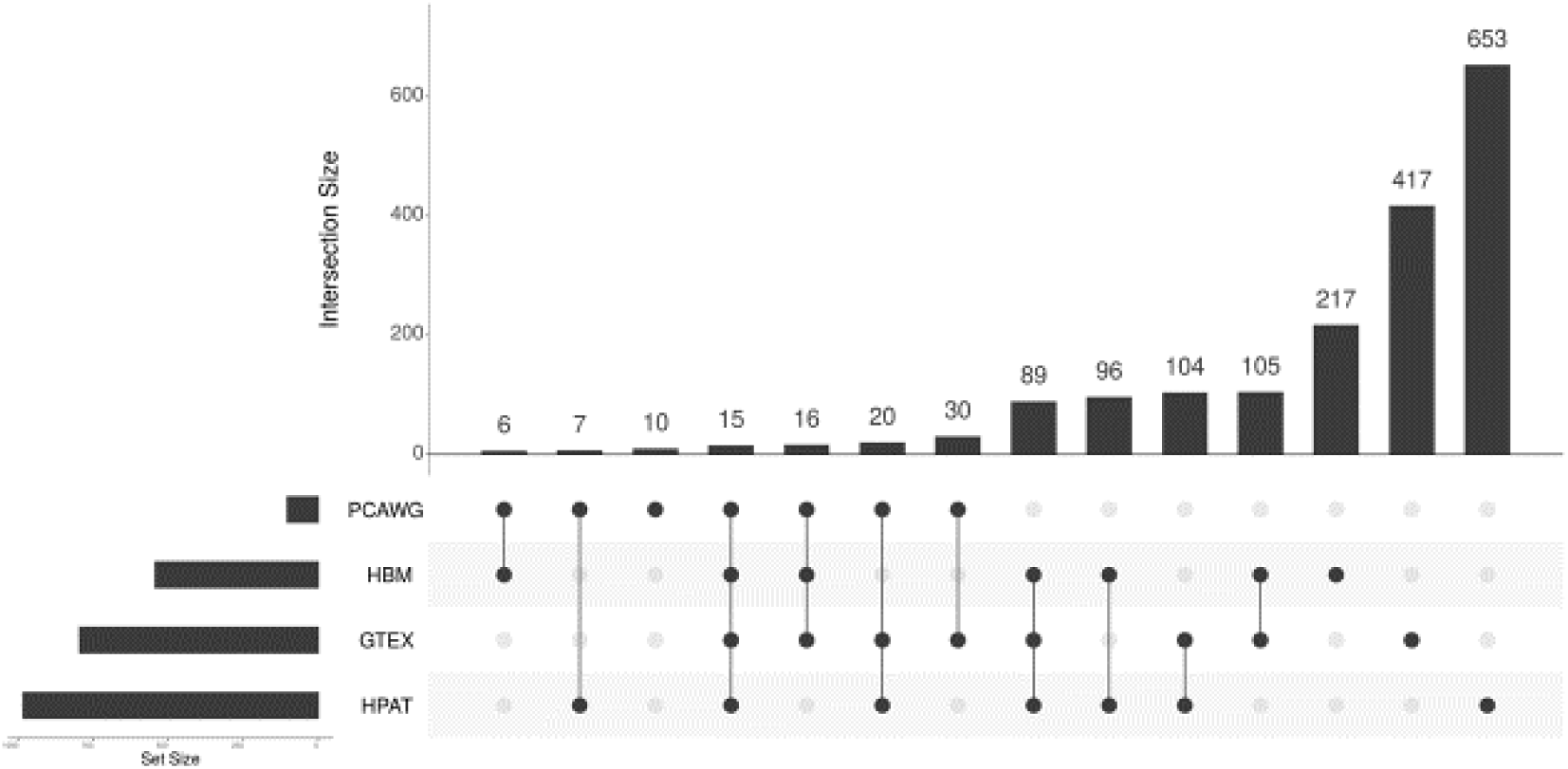
UpSetR [143] plot showing genes with a GC <0.2 that are variously shared and unique across the PCAWG, HBM, GTEX and HPA tissue data sets. The data underpinning this plot can be found it Supplementary Table S4

Sorting nexins are a group of cytoplasmic and membrane-associated proteins involved in the regulation of intracellular trafficking [116]. SNX3 has been reported to play a role in receptor recycling and formation of multivesicular bodies [117], and its dysregulation has been implicated in disorders of iron metabolism and the pathogenesis of some neurodegenerative diseases [118, 119].

The COX4I gene encodes the nuclear-encoded cytochrome c oxidase subunit 4 isoform 1, the terminal enzyme in the mitochondrial respiratory chain. Given the key role of the mitochondrial respiratory chain in all human cells (except red blood cells), stable expression of such a gene in all tissues may not be a surprising result. Increased RNA COX4I1 levels have been reported in sperm of an obese male rat model [120] and thus may play a role in obesity-related fertility problems, and reduced expression of this subunit leads to a reduction in mitochondrial respiration as well as sensitising cells to apoptosis [121].

The small number of genes shared between these data sets with a GC < 0.2 indicates that the data in these studies are more variable compared to cell lines alone. The cause of this variation may be due to the tissue data having been obtained from different subjects [122]. Moreover, tissues are themselves a mixture of cell types with varying levels of gene expression in each cell type [123], while cell lines are nominally clonal.

Our results suggest that in the case of RNA-seq tissue data sets, where gene expression tends to be more variable, an unbiased approach, using the Gini coefficient, may be more fruitful in the search for stably expressed genes with which to perform normalisation, than the other commonly used methods used until now [122, 124].

### RT-qPCR analysis of gene expression stability of some housekeeping genes in 10 cell lines

In order to illustrate the utility of the GC to find suitable housekeeping genes, we next chose to assess this experimentally by RT-qPCR using a small subset of candidate reference genes (40; top 32 genes from genes ordered by GC and expression value from [91], plus 8 of the most commonly used from the literature, including seven from [63] and one (RPL32) from [125][126], and 10 cell lines from a range of tissues (see Table 5 and 6). We first set a Cq value (which is inversely proportional to expression level) cut-off of 32, above which no expression is observed, and subsequently used the Cq values of genes in cell lines as a relative expression level (Cq cut off/Cq value of gene). Descriptive statistics of the expression of each gene in individual cell lines were then calculated. As a final step, the median expression value of each gene in individual cell lines was used to calculate descriptive statistics, including the GC, of gene expression across these cell lines. Figure 11 illustrates a KNIME workflow [127-129] that we wrote for this purpose. The raw data and descriptive statistics extracted are provided in Supplementary Tables S5 and S6 respectively, and the KMNIME analysis workflow in Supplementary File 1.

**Fig 11.**
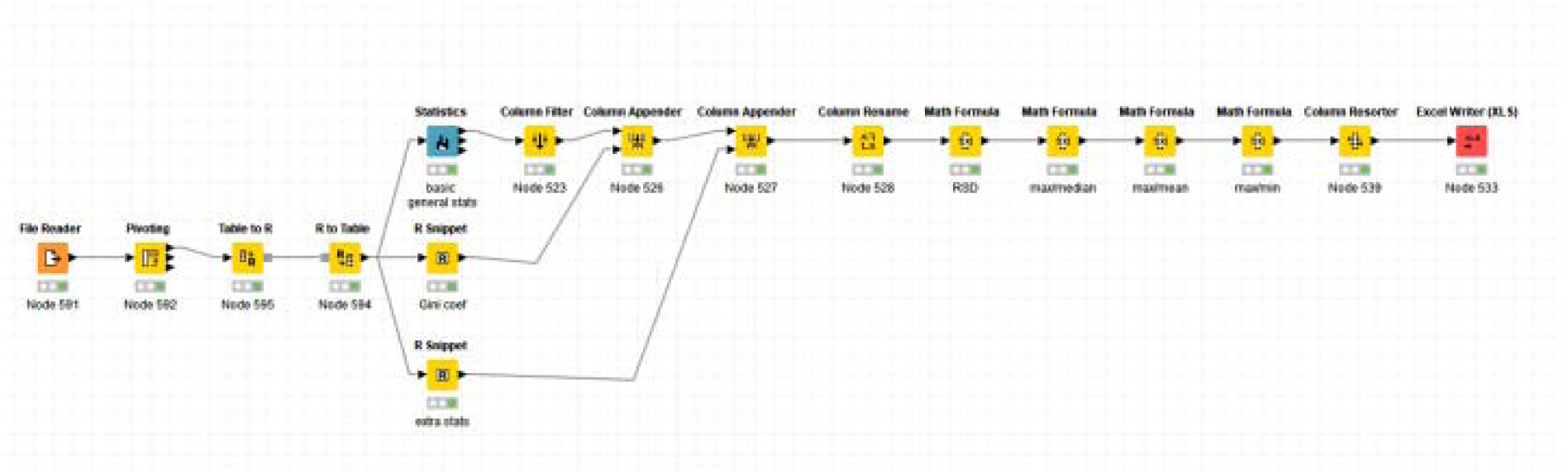
The KNIME workflow described here to calculate descriptive statistics and the gini coefficient from RT-qPCR data. This workflow can be adapted for use with large RNA-Seq Data sets.

Fig 12 uses RT-qPCR data to plot the GC of the candidate reference genes analysed here versus their relative median expression level. Three GiniGenes [91] (RBM45, TRNT1 and CNOT2) had very low and variable expression. Most of the other genes analysed showed low GC values with a range of (relative) expression values; the inset in Figure 12 shows genes with a GC < 0.2 including a mix of 35 genes: 26 GiniGenes and 6 housekeeping genes referenced by Vandesompele and colleagues [63], one referenced by Caracausi [130] and one by Lee et al [131]. Two of these GiniGenes, HNRNPK and PCBP1, which we also found to be stably expressed in the cell line data suggesting these may be potential stable housekeeping genes. As shown in Figure 13 and inset, the GC is well correlated with the % RSD.

**Fig 12.**
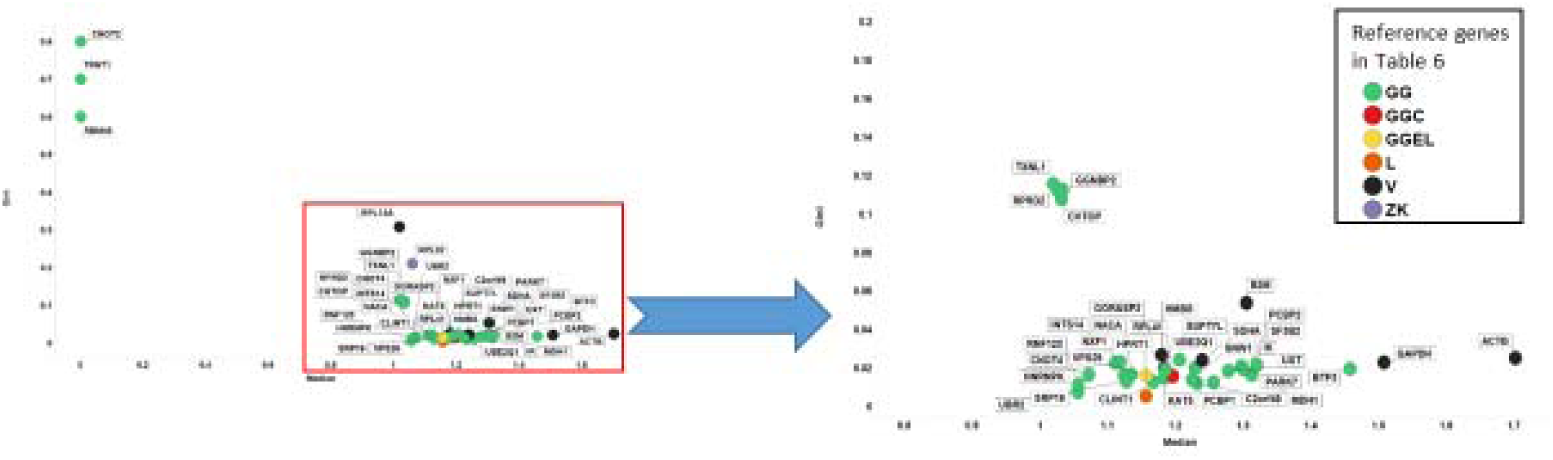
Gini coefficient and median expression levels of candidate reference genes assessed by RTqPCR. Left panel shows all genes considered in this study, with right panel showing genes with GC < 0.2. Colour coding: green, GeneGini reference genes; red, both GeneGini and Caracausi reference genes; yellow, GeneGini and Eisenberg and Levenson; orange, Lee, yellow; black, Vandesompele; purple, Zhang and Kriegova.

**Fig 13.**
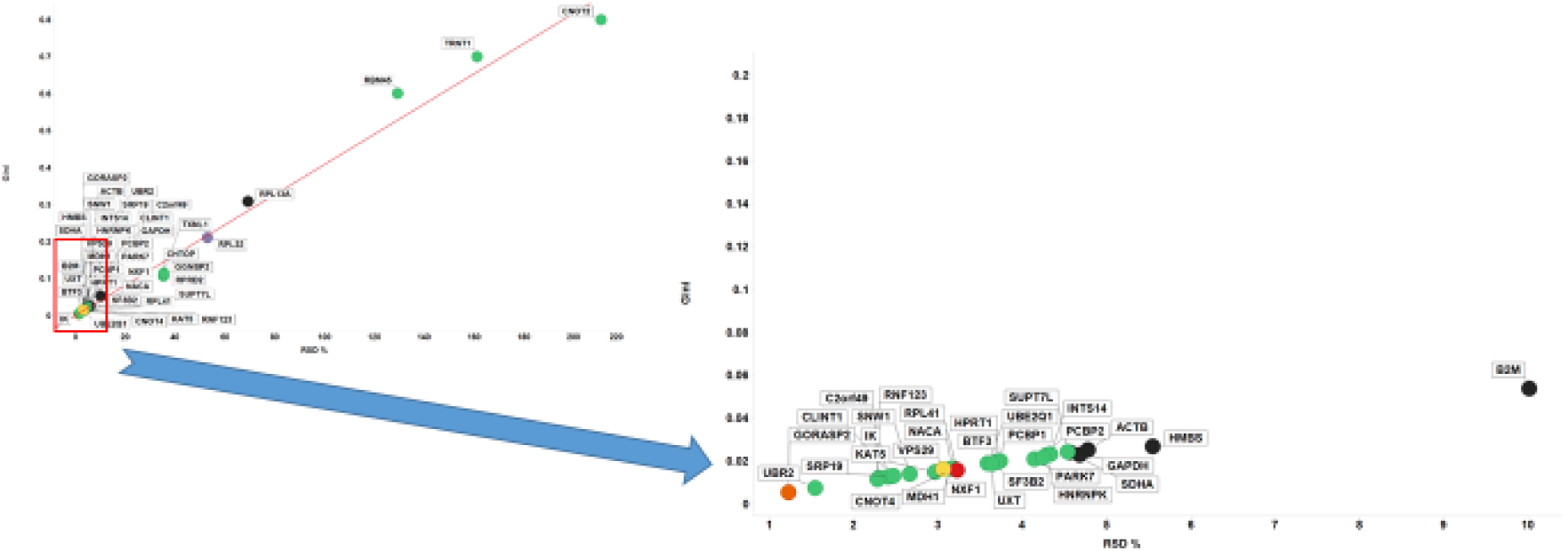
Robustness of the Gini coefficient in assessed experimentally by RT-qPCR using a small subset of proposed reference genes. Left panel shows Gini coefficient vs % RSD for all genes considered in this study, with right panel showing the same with genes with a GC < 0.2 and % RSD < 10. Line of best linear fit shown is y = 0.002 + 0.004x (r2=0.988). Shape coding as in Fig 12.

More importantly, the GC of our GiniGenes was particularly low (Fig 12). The low absolute magnitude reflected the fact that Cq value is based on a logarithmic scale. Various commonly used housekeeping genes (HPRT1, GAPDH, ACTB, SDHA, HMBS and B2M) displayed higher % RSDs and GC than other genes studied here in spite of their higher relative expression levels. This was also the case when inspecting the interquartile ratio against the GC of these (Figure S3).

The above results suggest that the GC is also applicable to RT-qPCR data, with GiniGenes having good potential (as novel “housekeeping” genes) for the normalisation of such data.

## Discussion

Reference genes are commonly used to normalise gene expression data, so as to account for bias resulting from both biological and technical variability, and to enable quantification of gene expression changes or differences in the system under study. It is generally considered that such reference genes should come from pathways that are required for general metabolism, using only one gene per ‘pathway’ to avoid co-regulation which might make the gene expressions look very stable.

Such reference genes are commonly referred to as ‘housekeeping’ genes (HKGs) because they are considered to participate in essential cellular functions, are ubiquitously expressed in all cells and tissue types, and their expression is considered to be stable [46-63]). A number of such genes have been proposed over the years, and genes such as GAPDH, ACTB, RPL13A, SDHA, B2M are frequently used in such studies [63]. However, the expression levels of these and other proposed HKGs have in fact been shown to vary widely between cells and tissues (e.g. [50, 59, 62, 64-76]) and their expression has also been reported to be affected by a number of factors relating to the experiment such as cell confluence [132], pathological, experimental and tissue specific conditions [133]. As highlighted by Huggett *et al*. [134], despite the reports of the potential variability of expression of ‘classic’ references genes such as GAPDH and ACTB, these are still used without mention of any validation processes. Our GiniGenes are selected as reference genes through different, data-driven, criteria.

Various tools have been developed to evaluate and screen reference genes from experimental datasets; these include geNorm [63], NormFinder [135], Best Keeper [136] and the comparative ΔCT finder [49]. RefFinder (http://leonxie.esy.es/RefFinder/#) and RefGenes can integrate these to enable a comparison and ranking of any tested candidate reference genes [137].

These tools assess expression stability of genes in different ways:

- geNorm determines gene stability through a stepwise exclusion or ranking process followed by averaging the geometric mean of the most stable genes from a chosen set. Python implementation: https://eleven.readthedocs.io/en/latest/
- BestKeeper also uses the geometric mean but using raw data rather than copy numbers. BestKeeper [136] can be used as an Excel-based tool. It can accommodate up to 10 housekeeping genes in up to 100 biological samples. Optimal HKGs are determined by pairwise correlation analysis of all pairs of candidate genes, and the geometric mean of the top ranking ones. http://www.gene-quantification.info
- NormFinder measures variation, and ranks potential reference genes between study groups. NormFinder [135] has an add-in for Microsoft Excel and is available as an R programme. It recommends analysis of 5-10 candidate genes and at least 8 samples per group. https://moma.dk/normfinder-software
- The comparative ΔCT finder requires no specialist programmes since this involves comparison of comparisons of ΔCTs between pairs of genes to find a set of genes that show least variability.
- RefGenes allows one to find genes that are stably expressed across tissue types and experimental conditions based on microarray data, and a comparison of results from geNorm, NormFinder and Best Keeper to find a set of reference genes. However, this is not a free service unless one searches for one gene at a time. Furthermore, the site for this tool is no longer available. Moreover, all these tools require the user to make a prior selection of such HKGs (introducing bias and potential errors) and most are cumbersome to understand and calculate.

We have here shown how via a simple calculation, the GC, we can find potential reference genes, and illustrated its utility in large-scale cell-line, tissue RNA-Seq data sets and RT-qPCR data. The expression of a number of classical HKGs from a number of carefully selected publications do in fact vary much more substantially between large RNA-Seq data sets, both for tissues and cell lines.

Whilst not all studies will involve large data sets such as those we have analysed here, the GC should also be of use for smaller-scale studies to select a subset of genes in a panel of cell lines or tissues relevant to the study in question.

Overall we find that (i) two of these genes, HNRNPK and PCBP1, seemed to be particularly robustly and stably expressed at reasonable levels in all cell lines studied, and (ii) a data-driven strategy based on the GC represents a useful and convenient method for normalisation in gene expression profiling and related studies.

## Methods

The datasets used are described and referenced below. The data, in transcripts per million (TPM) units were downloaded from the EBI expression atlas as a .tsv file. As previously [1], the Gini Index was calculated using the **ineq** package (Achim Zeileis (2014). ineq: Measuring Inequality, Concentration, and Poverty. R package version 0.2-13. https://CRAN.R-project.org/package=ineq) in **R** (https://www.R-project.org/). These calculations were incorporated into KNIME via KNIME’s R integration *R Snippet* node. A spreadsheet giving the extracted analyses is provided as supplementary tables (Tables S7 and S8).

### Cell lines and culture conditions

A panel of 10 cell lines were grown in appropriate growth media: K562, PNT2 and T24 in RPMI-1640 (Sigma, Cat No. R7509), Panc1 and HEK293 in DMEM (Sigma, Cat No. D1145), SH-SY5Y in 1:1 mixture of DMEM/F12 (Gibco, Cat No. 21041025), J82 and RT-112 in EMEM (Gibco, Cat No. 51200-038), 5637 in Hyclone McCoy’s (GE Healthcare, Cat No. SH30270.01) and PC3 in Ham’s F12 (Biowest, Cat No. L0135-500). All growth media were supplemented with 10 % fetal bovine serum (Sigma, Cat No. f4135) and 2 mM glutamine (Sigma, Cat No. G7513) without antibiotics. Cell cultures were maintained in T225 culture flasks (Star lab, CytoOne Cat No. CC7682-4225) kept in a 5% CO_2_ incubator at 37°C until 70-80 % confluent.

### Harvesting Cells for RNA Extraction

Cells from adherent cell lines were harvested by removing growth media and washing twice with 5 mL of pre-warmed phosphate buffered saline (PBS) (Sigma, Cat No. D8537), then incubated in 3 mL of 0.025% trypsin-EDTA solution (Sigma Cat No. T4049) for 2-5 min at 37 °C. At the end of incubation cells were resuspended in 5-7 mL of respective media when cells appeared detached to dilute trypsin treatment. The cell suspension was transferred to 15 mL centrifuge tubes and immediately centrifuged at 300 x g for 5 min. Suspended cell lines were centrifuged directly from cultures in 50 mL centrifuge tubes and washed with PBS as above. The cell pellets were resuspended in 10-15 mL media and cell count and viability was determined using a Nexcellom Cellometer Auto 1000 Cell Viability Counter (Nexcellom Bioscience) set for Trypan Blue membrane exclusion method. Cells with >95 % viability were used for downstream total RNA extraction.

### RNA Extraction

Total RNA was extracted from 2-5 × 10^6^ cells using the Qiagen RNeasy Mini Kit (Cat No. 74104) and DNAse treated using Turbo DNA-free kit (Invitrogen, Cat No. AM1907) according to the manufacturer’s instructions. Briefly, 1 X DNA buffer was added to the extracted RNA prior to adding 2U (1 µL) of DNAse enzyme. The reaction mixture was incubated at 37 °C for 30 min and inactivated for 2 min at room temperature using DNAse inactivating reagent. The mixture was centrifuged at 10,000 x g for 1.5 min and the RNA from the supernatant was transferred to a clean tube. The RNA concentration was determined using a NanoDrop^®^ ND-1000 spectrophotometer and further validated using an Agilent 2100 bio-analyser coupled with 2100 Expert software system. Only RNA samples with an RIN (RNA Integrity Number) between 9-10 were selected for cDNA synthesis.

### Reverse Transcription and cDNA Synthesis

1 µg of RNA was reverse transcribed into cDNA. Briefly, a 20 µL reaction was setup by adding 1 µL each of oligodT (50 µM, Invitrogen, cat No. 18418020) and dNTP mix (10 mM, Invitrogen, Cat No. 18427-013) followed by adding an appropriate volume for 1 µg of RNA. Nuclease free water (Ambion, Cat No. AM9937) was then added to make the volume up to 13 µL and incubated at 65 °C for 5 min then cooled on ice for 1min. To initiate transcription 4 µL of 5 X first strand buffer (Invitrogen, Cat No. 1889832) and 1 µL each of 0.1 M DTT (Invitrogen, Cat No. 1907572), RNaseOUT™ (Invitrogen, Recombinant RNase Inhibitor, Cat No. 1905432) and SuperScript™ III RT (200 units/µL, Invitrogen, Cat No. 1685475) reverse transcriptase enzyme were added, mixed gently then incubated at 50 °C for 60 min followed by inactivation at 70 °C for 15 min. The cDNA was diluted 1:100 to be used in RT-qPCR experiment.

### Validation of gene expression by geNorm

A set of candidate reference genes (40; top 32 genes from genes ordered by GC and expression value from [91], plus 8 of the most commonly used from the literature including seven from [63]). RNAseq data were selected for validation of stable gene expression using geNorm [63]. First, a typical qPCR protocol was prepared from a master mix for each gene to be tested per cell line in triplicate. This consisted of 10 µL/well made by adding 0.8 µL of nuclease free water (Ambion), 5 µL of LC480 SYBR Green I Master (2 X conc. Roche, Product No. 04887352001), 0.1 µL each of forward and reverse primers (20 µM) (for primer and amplicon sequences see Supplementary Table S9) and 4 µL of 1:100 diluted cDNA in a 384 well qPCR plate (Starlab Cat. No. E1042-9909-C). The no template controls (NTC) for each gene were produced by replacing cDNA with 4 µL of nuclease free water. Thermal cycling conditions used were: one cycle of 95 °C for 10 min followed by 40 cycles of 95 C for 10 sec and 60 °C for 30 sec. qPCR was performed using Roche LightCycler LC480 qPCR platform. The fluorescence signals were measured in real time during amplification cycle (Cq) and also during temperature transition for melt curve analysis.

The mean Cq values were converted into relative values for a gene across all cell lines using ΔCq method [142]. Briefly, the lowest Cq value in a panel of cell lines for a gene was subtracted from all the values in that panel using the equation: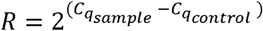, where 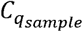 is the mean Cq value obtained for a gene in each of the cell lines and 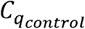 is the lowest Cq value in that panel. The relative values for each gene in a panel were then obtained by applying 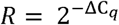. These relative values were applied in geNorm Visual Basic applet for Microsoft Excel^®^ [63] that determines the most stable reference genes from a set of genes in a given panel of cell lines.

### Validation of gene expression using the Gini coefficient

To the raw RT-qPCR data a Cq value (which is inversely proportional to expression level) cut-off of 32 was set, above which no expression is observed. The Cq values of genes in cell lines were subsequently converted to a relative expression level (Cq cut off/Cq value of gene). Descriptive statistics of the expression of each gene in individual cell lines were then calculated. As a final step, the median expression value of each gene in individual cell lines was used to calculate descriptive statistics, including the GC, of gene expression across these cell lines. Figure 11 illustrates a KNIME workflow [127-129] for this purpose. The raw data and descriptive statistics extracted are provided in Supplementary Tables S5 and S6 respectively, and the KMNIME analysis workflow in Supplementary File 1.

## Declarations

### Ethics approval and consent to participate

Not applicable.

### Consent for publication

The PCAWG data is under embargo until the WGS pan-cancer consortium publishes its marker paper or until July 25, 2019, whichever is earlier. Methodology papers may be published prior to this embargo, with agreement from the full scientific working group. We have been in email contact with who asked that we advise the editor to wait until the July 25 embargo lift.

### Availability of data and materials

All data generated or analysed during this study are included in this published article (and its supplementary information files). The original datasets used are referenced throughout and are summarised in Table 2.

### Authors’ contributions

D.B.K. highlighted the utility of the GC as shown in [1]. M.W.M. adapted the Gini method and analyses workflows developed by S.O. from [1] and performed most of the analyses that were done using KNIME. P.J.D. contributed in particular to the analysis of the housekeeping genes. F.M. performed the RT-qPCR analyses. All authors contributed to the writing and approval of the manuscript.

## Acknowledgements

All authors thank the BBSRC (grant BB/P009042/1) and the Novo Nordisk Foundation (grant NNF10CC1016517) for financial support. for financial support.

## Legends to figures

**Supplementary Fig S1.**
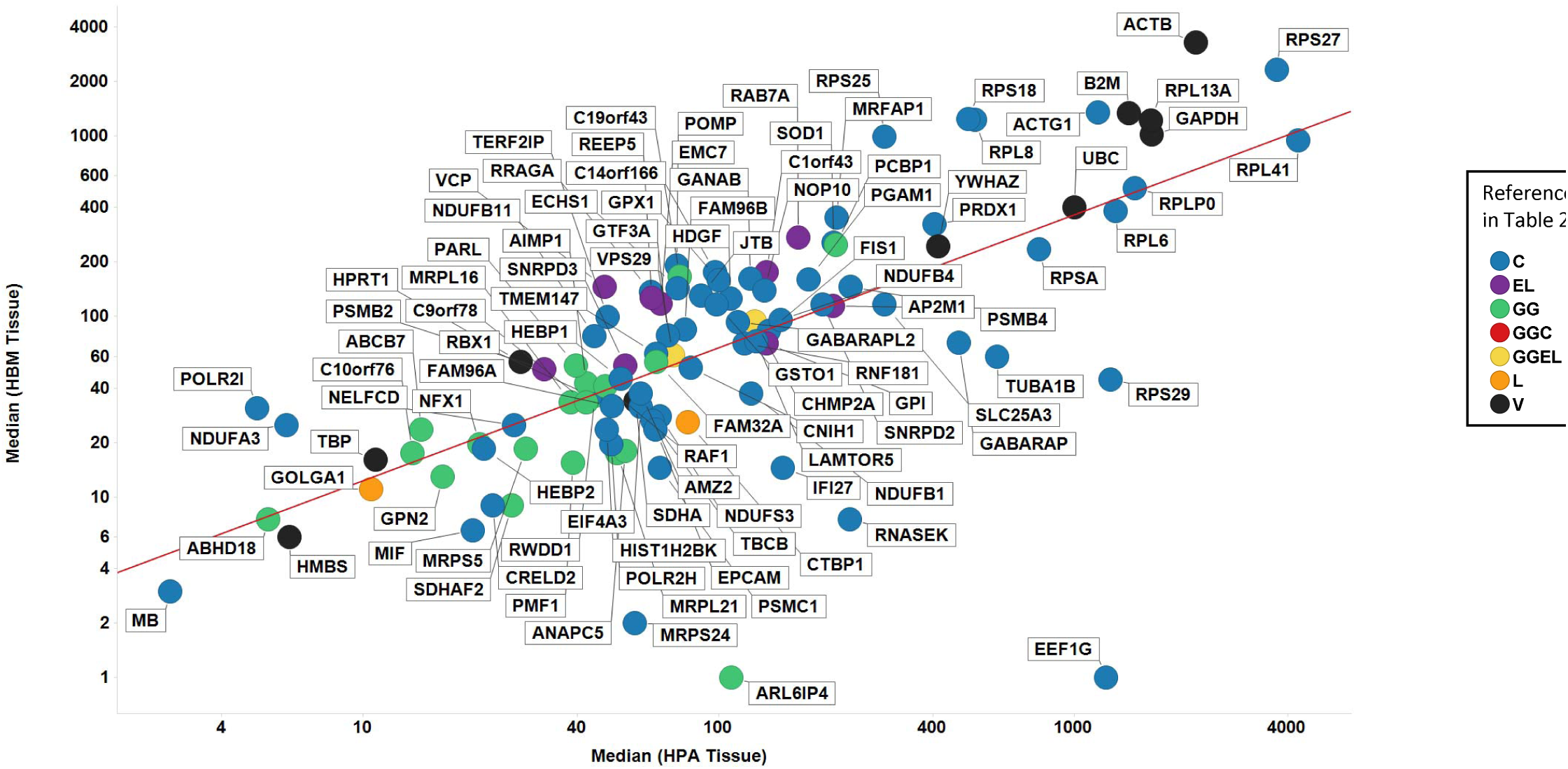
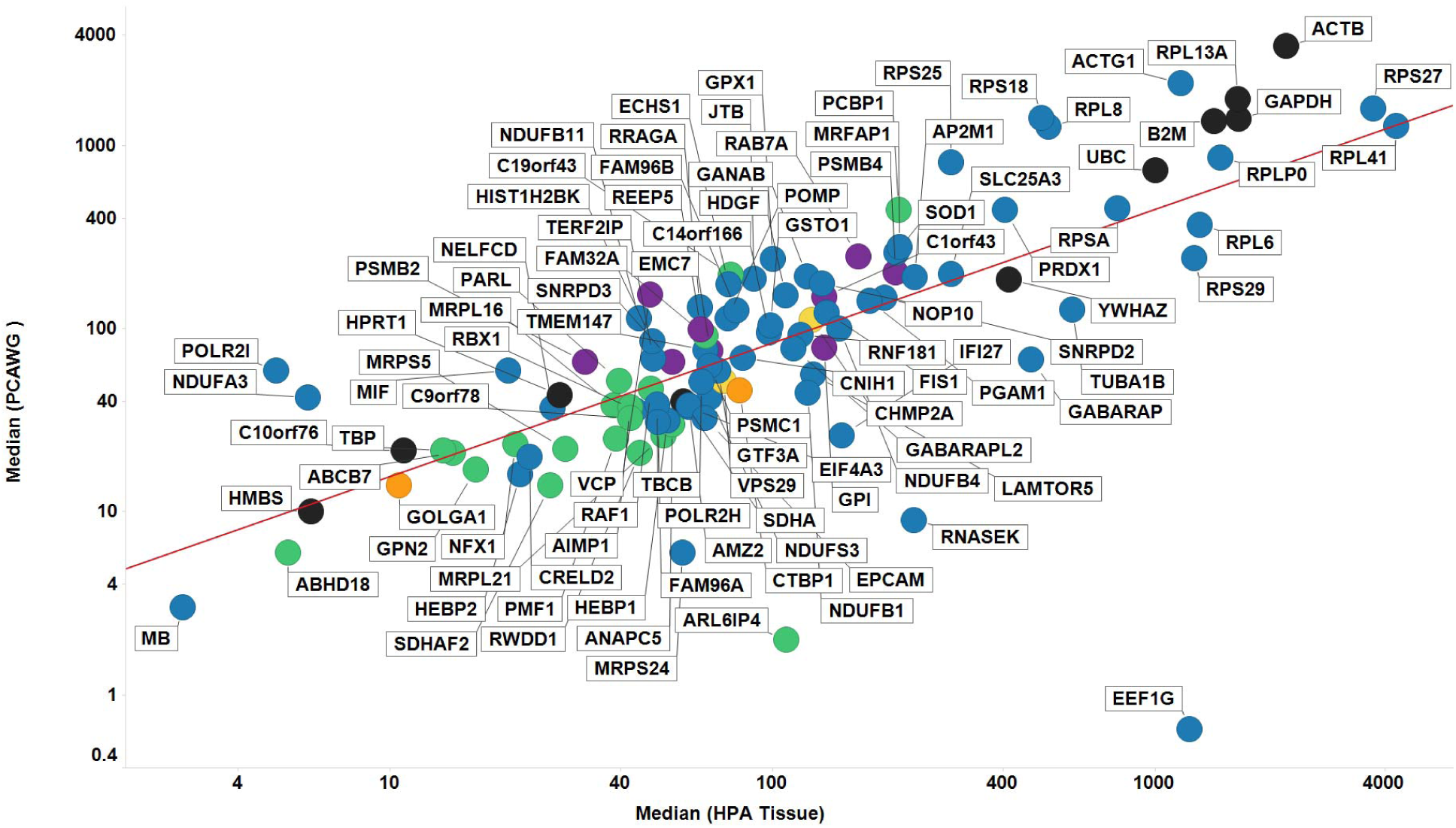
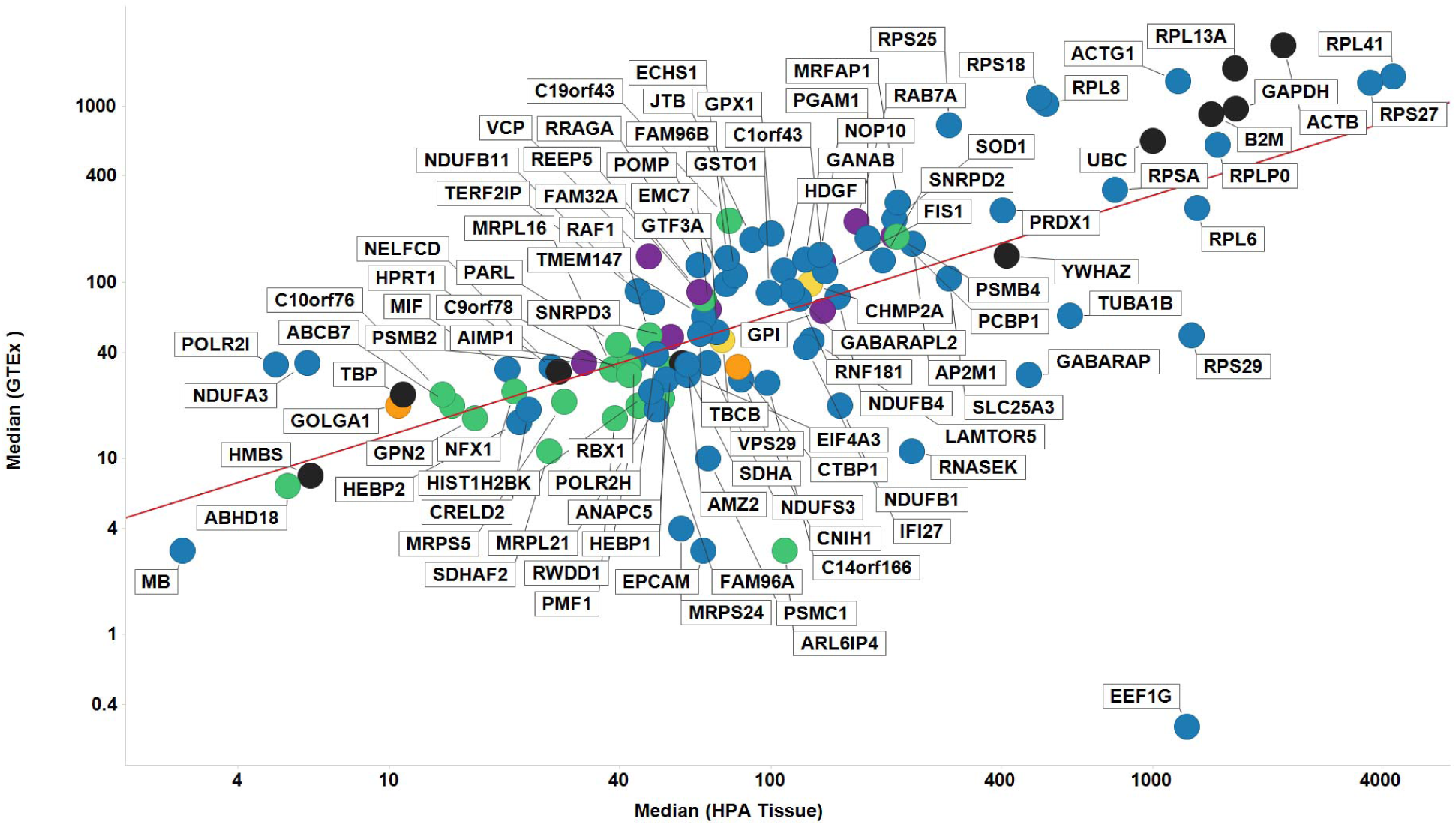
Comparison of median expression levels of proposed reference genes between tissue datasets. A. HBM vs HPA tissue datasets. Line of best linear fit (in log space) shown is log_10_y = 0.35 + (0.74 log_10_ (x)) (r2=0.472). B. PCAWG vs HPA tissue dataset. Line of best linear fit (in log space) shown is log_10_y = 0.46 + (0.73 log_10_ (x)) (r2=0.500). C. GTEx vs HPA Tissue. Line of best linear fit (in log space) shown is log_10_y = 0.45 + (0.68 log_10_ (x)) (r2=0.429). Colour coding: blue, Caracausi reference genes; purple, Eisenberg & Levenson; green, GeneGini; yellow, both GeneGini and Eisenberg and Levenson; orange, Lee; black, Vandesompele.

**Supplementary Fig S2.**
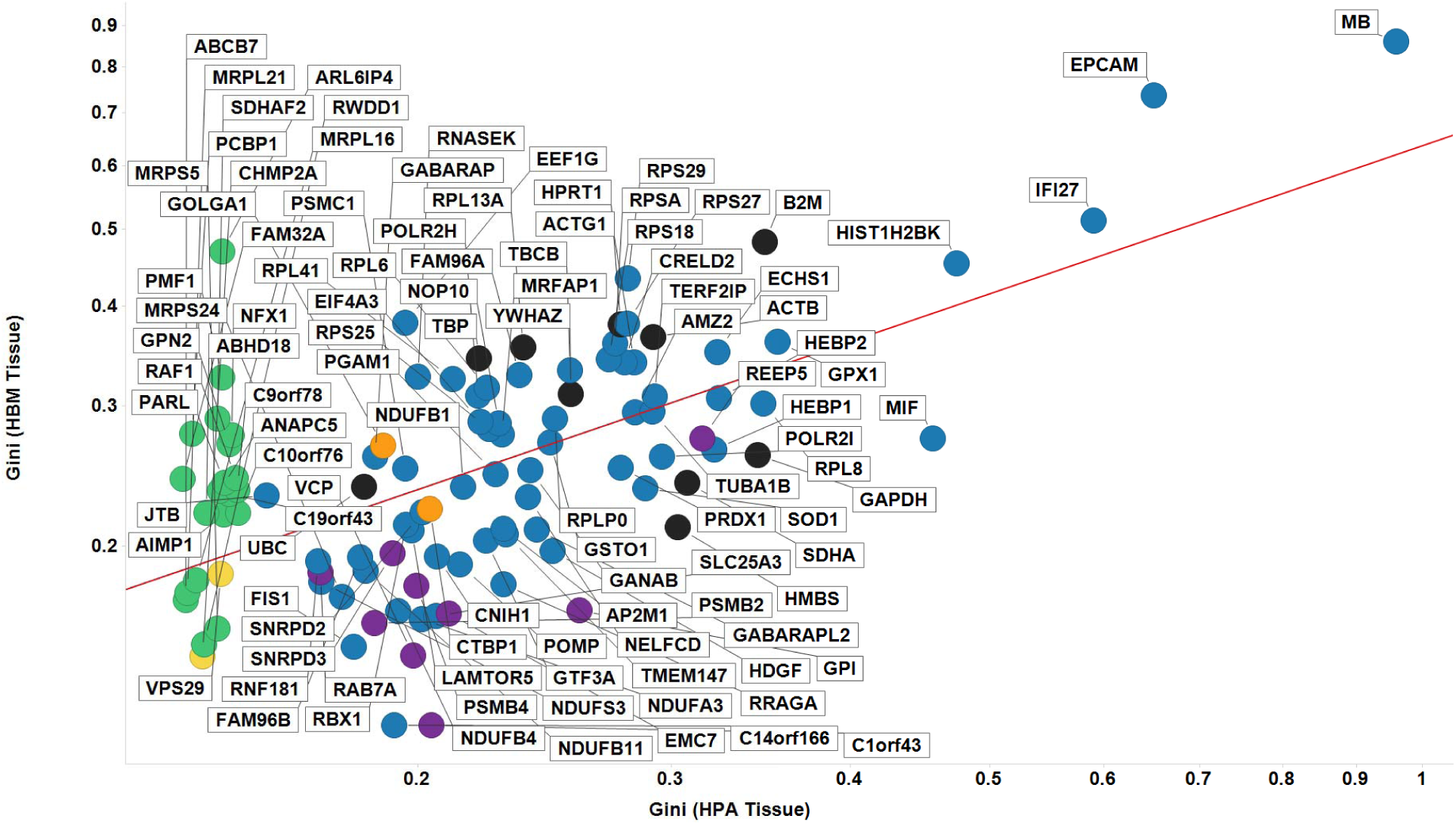
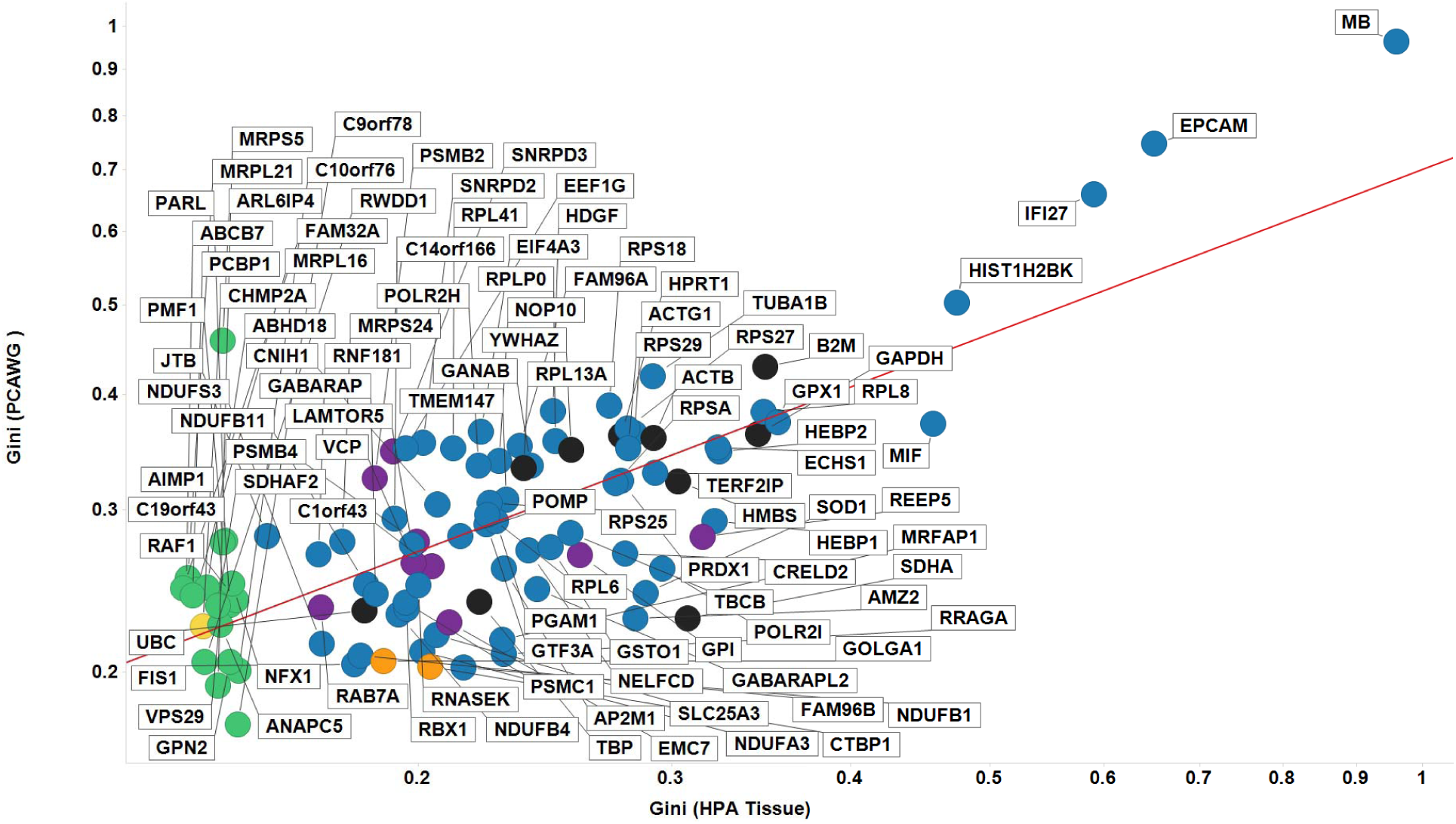
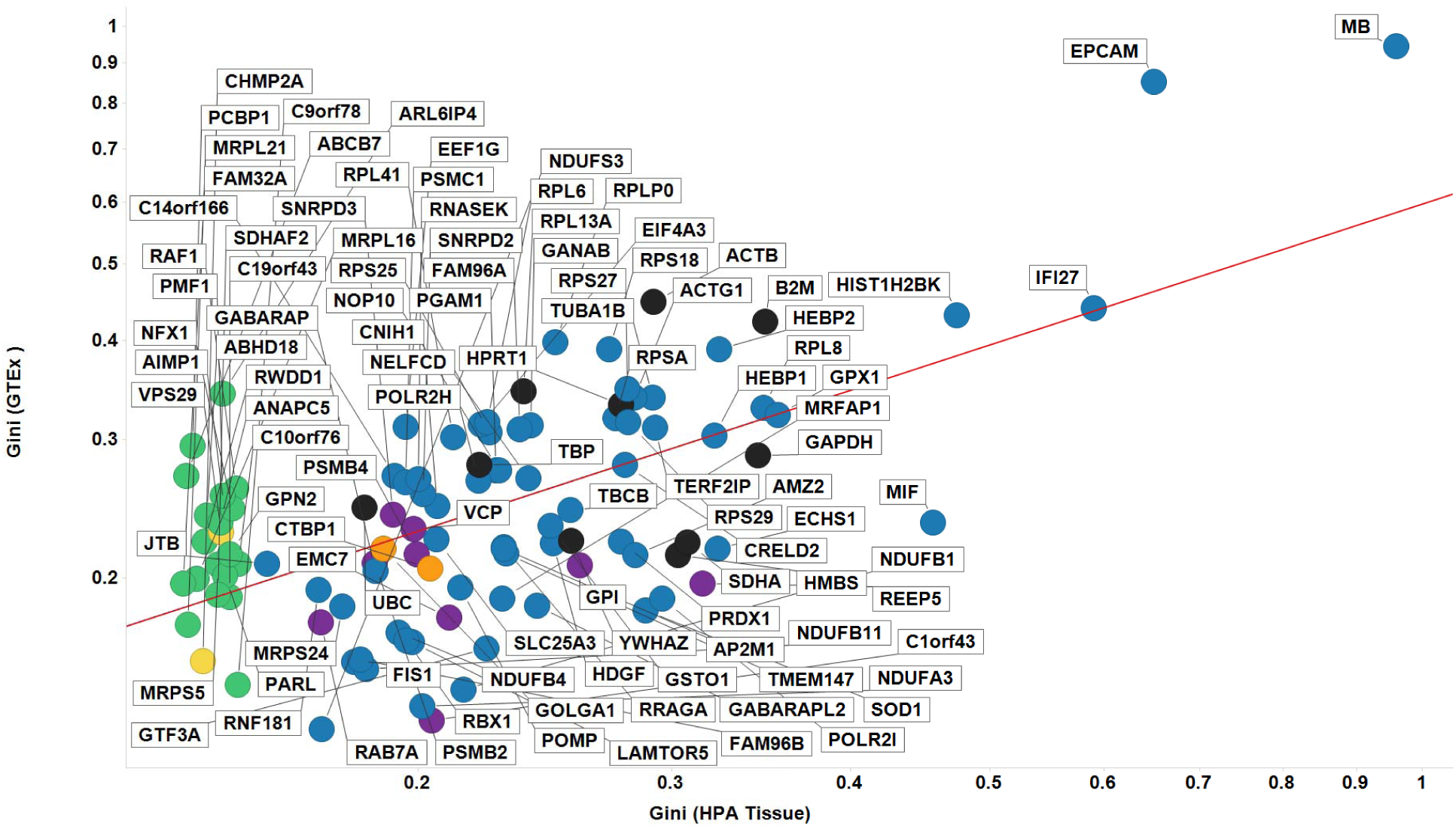
Comparison of Gini coefficient of proposed reference genes between tissue datasets. A. HBM vs HPA tissue datasets. Line of best linear fit (in log space) shown is log10y = −0.20 + (0.62 log_10_(x)) (r2=0.392). B. PCAWG vs HPA tissue dataset. Line of best linear fit (in log space) shown is log_10_y = −0.15 + (0.59 log_10_ (x)) (r2=0.560). **C.** GTEx vs HPA Tissue. Line of best linear fit (in log space) shown is log_10_y = 0.22 + (0.59 log_10_ (x)) (r2=0.388). Colour coding as in Fig S1.

**Fig S3.**
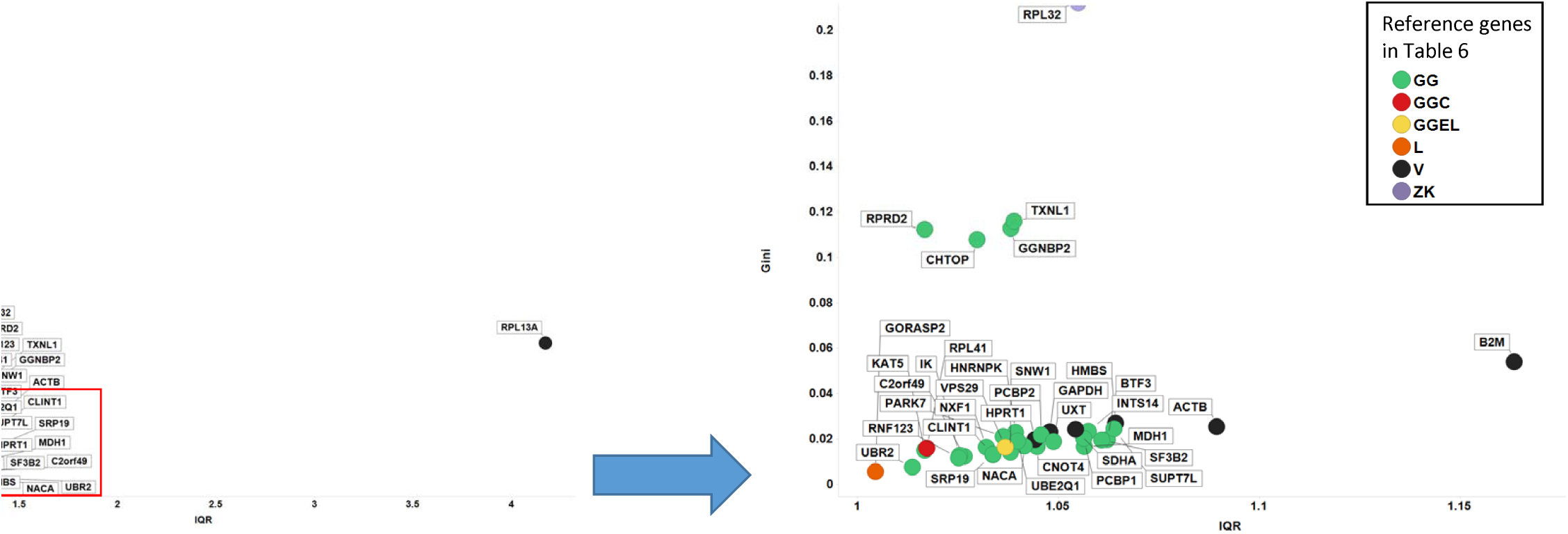
Robustness of the Gini coefficient assessed experimentally by RT-qPCR using a small subset of proposed reference genes illustrated with Gini coefficient vs IQR. Left panel shows all 40 genes in Table 6, with right panel showing genes with a GC < 0.2. Colour coding: green, GeneGini reference genes; red, both GeneGini and Caracausi reference genes; yellow, GeneGini and Eisenberg and Levenson; orange, Lee, yellow; black, Vandesompele; purple, Zhang and Kriegova.

## List of tables

Table 3. Descriptive statistics of 13 genes common across cell-line data sets with GC < 0.2. In addition, the protein name, as well as UniProt ID and function are shown. S/A/O refers to SLC, ABC or Other respectively.

Table 4. Descriptive statistics of 15 common genes across tissue data sets with a GC < 0.2. In addition, the protein name, as well as UniProt ID and function are shown.

Table 5. Details of human cell lines used for the assessment of expression of candidate reference genes by RT-qPCR.

Table 6. Candidate reference genes used to assess expression stability experimentally by RT-qPCR. Included are gene name and UniProt ID, Gini coefficient as calculated using the HPA cell-line data set. S/A/O refers to SLC, ABC or Other respectively.

Supplementary Table S1. Descriptive statistics of 115 common genes across cell-line datasets. S/A/O refers to SLC, ABC or Other respectively.

Supplementary Table S2. Descriptive statistics and UniProt names and IDs of proposed stable reference genes from Table 1 in tissue datasets. S/A/O refers to SLC, ABC or Other respectively.

Supplementary Table S3. Descriptive statistics of common and unique genes across tissue data sets with a GC ≤ 0.2, including gene names and functions. S/A/O refers to SLC, ABC or Other respectively.

Supplementary Table S4. Data underpinning UpSetR [143] plot in Figure 10 showing genes with a GC <0.2 that are variously shared and unique across the PCAWG, HBM, GTEX and HPA tissue data sets.

Supplementary Table S5. Raw expression data for candidate reference genes in human cell lines by RT-qPCR.

Supplementary Table S6. Descriptive statistics and Gini coefficient data for candidate reference genes in human cell lines by RT-qPCR.

Supplementary Table S7. Extracted analyses of cell-line RNA-Seq data sets referenced in Table 2. S/A/O refers to SLC, ABC or Other respectively.

Supplementary Table S8. Extracted analyses of tissue RNA-Seq data sets referenced in Table 2. S/A/O refers to SLC, ABC or Other respectively.

Supplementary Table S9. Primer and amplicon sequences of candidate reference genes used to assess expression stability experimentally by RT-qPCR. Included are the Gini coefficient and median expression level as found in the HPA cell-line data set. S/A/O refers to SLC, ABC or Other respectively.

## Supplementary Files

Supplementary File 1. KNIME workflow [127-129] that we have written to calculate descriptive statistics, including the GC, of gene expression across cell lines to assess of expression stability of candidate reference genes by RT-qPCR.

